# Active membrane deformations of a minimal synthetic cell

**DOI:** 10.1101/2023.12.18.571643

**Authors:** Alfredo Sciortino, Hammad A. Faizi, Sarvesh Uplap, Layne Frechette, Matthew S. E. Peterson, Petia Vlahovska, Aparna Baskaran, Michael F. Hagan, Andreas R. Bausch

**Affiliations:** Lehrstuhl für Biophysik (E27), Physik Department, Technische Universität München, Garching bei München, 85748 Germany; Center for Protein Assemblies, Garching bei München, 85748 Germany; Department of Mechanical Engineering, Northwestern University, Evanston, IL 60208, USA; Martin A. Fisher School of Physics, Brandeis University, Waltham, MA, 02453, United States; Department of Engineering Sciences and Applied Mathematics, Northwestern University, 60208, USA

**Author notes:** These authors contributed equally (Alfredo Sciortino, Hammad A. Faizi).

## Abstract

Biological cells exhibit the remarkable ability to adapt their shape in response to their environment, a phenomenon that hinges on the intricate interplay between their deformable membrane and the underlying activity of their cytoskeleton. Yet, the precise physical mechanisms of this coupling remain mostly elusive. Here, we introduce a synthetic cell model, comprised of an active cytoskeletal network of microtubules, crosslinkers and molecular motors encapsulated inside giant vesicles. Remarkably, these active vesicles exhibit large shape fluctuations and life-like morphing abilities. Active forces from the encapsulated cytoskeleton give rise to large-scale traveling membrane deformations. Quantitative analysis of membrane and microtubule fluctuations shows how the intricate coupling of confinement, membrane material properties and cytoskeletal forces yields fluctuation spectra whose characteristic scales in space and time are distinctly different from passive vesicles. We demonstrate how activity leads to uneven probability fluxes between fluctuation modes and hence sets the temporal scale of membrane fluctuations. Using simulations and theoretical modelling, we extend the classical approach to membrane fluctuations to active cytoskeleton-driven vesicles, highlighting the effect of correlated activity on the dynamics of membrane deformations and paving the way for quantitative descriptions of the shape-morphing ability typical of living systems.

---

Rather than being merely passive containers, cell membranes actively respond to and steer cellular activity, enabling a myriad of biological functions such as cell crawling, cell division and cytoplasmic streaming [1–8]. In order to accomplish these activities, cells must have the ability to dramatically change their shape. Many of these processes arise from a tight coupling between lipid membrane fluctuations, providing the necessary flexibility, and the underlying cytoskeleton, which is in a highly non-equilibrium state and provides the necessary forces and directionality to induce deformations. The seminal discovery of “membrane flickering” in red blood cells [9, 10] revealed how fluctuation analysis is essential to understand membrane driven processes both *in vivo* and *in vitro*. One general result at equilibrium is that the temporal relaxation of membrane fluctuations is tightly bound to its spatial correlations, as dictated by Onsager regression and the resulting fluctuation-dissipationb theorem [11–14]. This dependence enables extracting mechanical properties from both spatial and temporal measurements of membrane dynamics. However, in cells, the cytoskeletal activity modifies both the spatial and the temporal behaviors of membrane deformations [15], potentially breaking the interdependence of spatial and temporal fluctuations dictated by thermodynamics [14, 16–19]. Yet, in living systems, it is challenging to simultaneously measure the dynamics of the membrane and that of the cytoskeleton imparting forces on it. This makes it unfeasible to explicitly connect cytoskeletal activity to the resulting fluctuations.

Giant unilamellar vesicles (GUVs) are a powerful tool to investigate in a controlled minimal system how membrane deformations are affected by activity [20–24]. However, despite recent advancements [24–36] the realization of a minimal experimental system exhibiting the shape-morphing ability of cells due to an active, complex, three-dimensional cytoskeleton is still lacking. Here we address this issue by encapsulating a reconstituted cytoskeleton composed of a crosslinked active microtubule 3D network inside a deformable GUV, and analysing the resulting fluctuations and shape deformations.

## Vesicle shapes induced by active bundles

Our experimental model of the cytoskeleton consists of an active gel composed of microtubules and molecular motors [26, 37–41], which we encapsulate using the cDICE technique[25, 42]. Microscopically, the system is composed of short microtubules (MTs, at a concentration *c*_*MT*_), kinesin tetramers (*c*_*K*_) and the protein anillin (*c*_*A*_) as a crosslinker (Fig. S1.A). Powered by molecular motors, this active cytoskeleton self-organizes into a loosely connected, isotropic 3D active network of long bundles (≈ 100 *μm*) that continuously extend and buckle due to kinesin-induced stress (Fig. 1.a).

**Figure 1.**
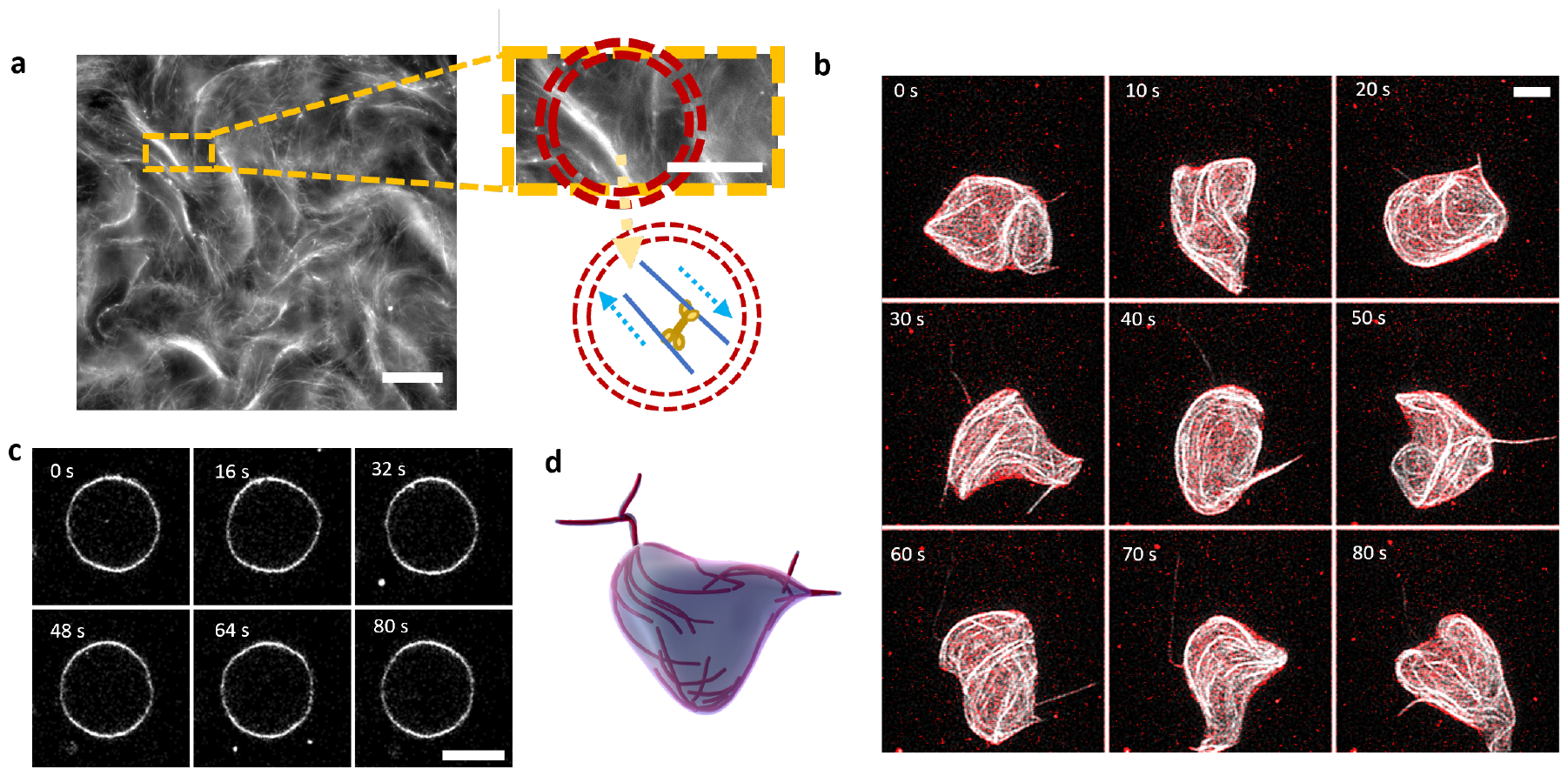
GUVs containing a minimal cytoskeleton exhibit large shape fluctuations: **a**, Confocal snapshot of the active MT network in bulk. Short fluorescently labelled MTs, in the presence of crosslinkers, self-organize into extensile active bundles, propelled by kinesin motors (see schematics on the right). Scale bar is 100 *μm*. Right, top) Enlargement of the extensile bundles. A circle having a size comparable to the GUVs we produce (*R*_0_ ≈ 25 *μm*) is superimposed in red for reference. Scale bar is 50 *μm*. Right, bottom) Schematics of the encapsulated experimental system. Inside the GUV (red circle), microtubules of opposite polarities (blue arrows) are extended by kinesin motors, resulting in active bundles confined inside the GUV. **b**, Confocal projections of a GUV (membrane in red) containing an active microtubule network (MTs in white). The GUV deforms and changes shape with a timescale of the order of a few seconds. Scale bar is 20 *μm*. **c**, A passive vesicle fluctuating with the same time interval between frames as a reference. Scale bar is 20 *μm*. **d**, Artist’s depiction of the shape-morphing GUVs resulting from the encapsulation of active bundles.

When encapsulated inside GUVs (mean radius *R*_0_ ≈ 25*μm*), the active microtubule network spans the whole vesicle volume, in stark contrast to previous systems in which a dense nematic network was confined to a 2D layer at the membrane [26]. The active forces exerted by the microtubule gel on the membrane induce large shape deformations of the vesicle (Fig. 1.b, Movie 1). The GUV continuously undergoes dramatic morphological changes, with a timescale of ≈ 10 *s*, as extracted by the correlation function of membrane deformations (Fig. S4.C). Active deformations are clearly different in magnitude and dynamics from those present at equilibrium (Fig. 1.c). They are observed for a wide range of concentrations of microtubules, kinesin and anillin (Fig. S1.B). These enhanced deformations appear to be tightly correlated with the organization of the microtubule network inside the GUV (Figs. 1.d, Movie 2). Moreover, vesicles never settle into a definite shape as observed in previous systems [20, 22, 24], but rather continuously oscillate around a spherical geometry.

## Fluctuation spectroscopy reveals enhanced out-of-equilibrium fluctuations

To gain further insight into the spatio-temporal dynamics of active GUVs, we resort to flicker spectroscopy. We take long, high-framerate (30 to 4 frames per second, Movie 3) images of the equatorial plane of a deforming GUV and then decompose its contour into Fourier modes, denoted by the letter *q* (Fig. 2.e, SI), as

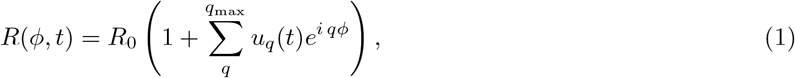

where *q*_max_ ≈ 15 is fixed by the image resolution. Each mode *u*_*q*_ represents the magnitude of equatorial deformations of wavelength ≈ *R/q*.

**Figure 2.**
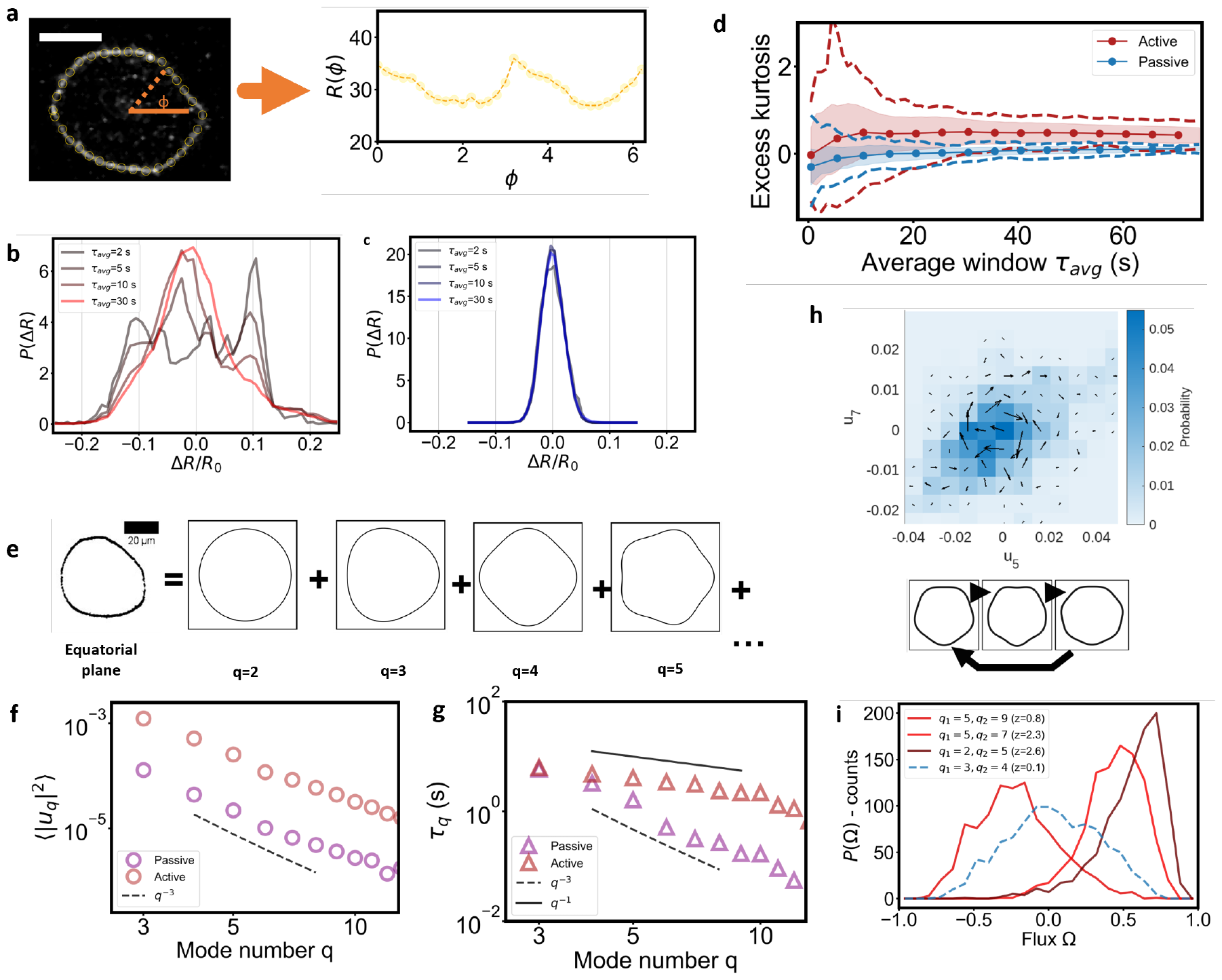
Membrane deformations of active vesicles are out-of-equilibrium: **a**, To analyse deformations, the equatorial contour of a fluctuating vesicle is tracked to obtain its description in polar coordinates *R*(*ϕ, t*). The extracted contour is marked by orange dots and then shown as a plot on the right. Scale bar is 25 *μm*.**b, c**, Histograms of radial deformations ∆*R* for an active GUV (**b**), compared to a passive one (**c**). The contours are sampled for different times (*τ*_*avg*_ =2, 5, 10, 30 s) from movies of GUVs. Active deformations have peculiar distributions at short time scales indicating correlated dynamics at short time scales. Passive GUVs exhibit a gaussian-like distribution at all time-scales, as expected from thermal noise. **d**, Excess kurtosis of the distribution of radial deformations as a function of the sampling time *τ*_*avg*_. By extracting random trajectories of given duration *τ*_*avg*_ we can obtain the mean kurtosis (circles), its standard deviation (shaded area), and the minimum and maximum kurtosis (dashed lines) at a given sampling time. While passive distributions always have an excess kurtosis close to 0, comparable with that of a Normal distribution, and a small spread around it, active ones at short time scales have a larger spread of kurtosis and can reach more extreme values. Only at longer times is a more symmetric distribution reached. **e**, Schematic of the decomposition of the contour into a sum of Fourier modes, labelled with *q*, whose fluctuations can then be separately analysed. **f**, Fluctuation spectrum as a function of the Fourier mode *q* for a passive GUV (purple) and an active GUV (red). Fluctuations of the active GUV are higher in magnitude and decay with a scaling exponent of ≈ 3. The dashed line indicates a *q*^−3^ scaling. **g**, Correlation time at each mode *q* for a passive (purple) and an active (red) vesicle, as obtained from the correlation functions 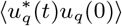. While the passive vesicle exhibits the expected *q*^−3^ scaling (dashed line) the active one has a different one (solid line indicates *q*^−1^). **h**, We explicitly measure how detailed balance is broken by measuring the probability flux Ω between different Fourier modes *q*_1_ and *q*_2_. The schematic on the bottom illustrates the transition between shapes dominated by the above Fourier modes. **i**, Histogram of *P* (Ω) showing net fluxes between pairs of different modes, which is a signature of out-of-equilibrium activity. However, we find that not all pairs of modes show a net flux. The dashed blue line shows instead an example from a passive vesicle showing no net flux. For each dataset we compute the z-score as the mean divided by the standard deviation (see SI), indicating how far the mean is from Ω = 0.

For passive vesicles with bending rigidity *k* and tension *σ*, the power spectrum of the Fourier coefficients *u*_*q*_ is expected to scale as

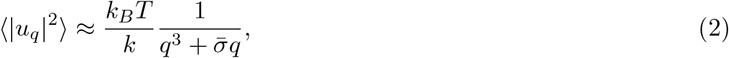

where *k*_*B*_*T* is the thermal energy and 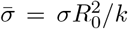 is a normalized tension. Note that this equation for the equatorial fluctuations of a GUV is the equivalent of the classical Canham-Helfrich model of a planar membrane [11–13] projected on a single dimension. We recover for passive vesicles the classical regimes of bending 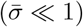 (Fig 2.f, purple) and tension dominated fluctuations 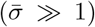 (Fig. S2.A). From this data we extract for passive GUVs made of Egg phosphatidylcholine (EggPC) the bending regidity of *κ*_pass_ = (13.4 ± 2.5) *k*_*B*_*T* and a membrane tension *σ* ≈ 10^−7^ *N/m* consistent with the literature [43, 44], confirming that the cDICE approach to prepare GUVs does not affect their mechanical properties. Vesicles encapsulating passive microtubule networks in the absence of ATP, or also in the presence of short microtubules without motors and crosslinkers, do not show any peculiar fluctuation spectra, and their dynamics resemble those observed for bare vesicles (Fig. S2.B).

In contrast, for actively deforming GUVs we observe fluctuations that are roughly one order of magnitude above the passive reference (Fig. 2.f, red) at all modes (from now on, all graphs refer to the same vesicle containing 0.8 mg/ml MTs, *c*_*A*_ = 1.5 *μ*M and *c*_*K*_ = 120 nM; further examples shown in S2.C). The fluctuation spectra of active GUVs decay over *q* similarly to the observed passive bending-dominated case ⟨|*u*_*q*_|^2^⟩ ∼ *q*^−3^. The increase in magnitude indicates that, in the presence of activity, thermal excitations of the bending modes of the vesicle are negligible compared to the deformations driven by active forces [20, 21, 45].

We then turn to the temporal behavior of the deformations. The time correlation function 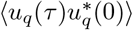 of a purely passive GUV is expected to decay as an exponential [13] with a decay time *τ*_*q*_, which in turn scales exactly like the spatial spectrum as 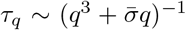 (Fig. 2.g). For active GUVs, however, we observe a 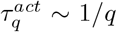 scaling at low modes, which does not match the scaling of spatial fluctuations. This indicates that activity strongly affects the timescales of the membrane fluctuations, and thus defies the relationship between fluctuations and response anticipated by the fluctuation-dissipation theorem, that would impose the same scaling between spatial fluctuations and their temporal decay. Indeed, using previously established methods [46, 47], we confirm that membrane fluctuations do break detailed balance. Briefly, using the amplitude of Fourier modes as a proxy for the GUVs’ microscopic configurations over time, we find the presence of net probability fluxes Ω in the transitions between different modes, which are instead expected to vanish at equilibrium due to detailed balance (Fig. 2.h, Fig. S3). Intriguingly, we find the presence of a statistically significant net flux only between some couples of modes, indicating that the active force couples preferentially with deformations of specific wavelengths (Fig. 2.i). This confirms that the membrane is behaving as an out-of-equilibrium system and that its temporal dynamics can be informative about the microscopic details of activity-induced deformations.

## Microtubules set the correlation time of membrane fluctuations

Since the membrane fluctuations are induced by the contained minimal cytoskeleton, we now focus on properties of the MTs inside the GUV, taking advantage of our ability to image them at the same time as the membrane.

Because the bundles’ length (≈ 100 *μm*) is comparable with the GUVs’ diameter (≈ 50 *μm*), not only does the active fluid affect the membrane fluctuations, but, in turn, the MT organization is altered by the confinement of the membrane. This cross-talk between membrane and cytoskeletal activity leads to a complex, three dimensional organization of the bundles (Fig. 3.a, Movie 4).

**Figure 3.**
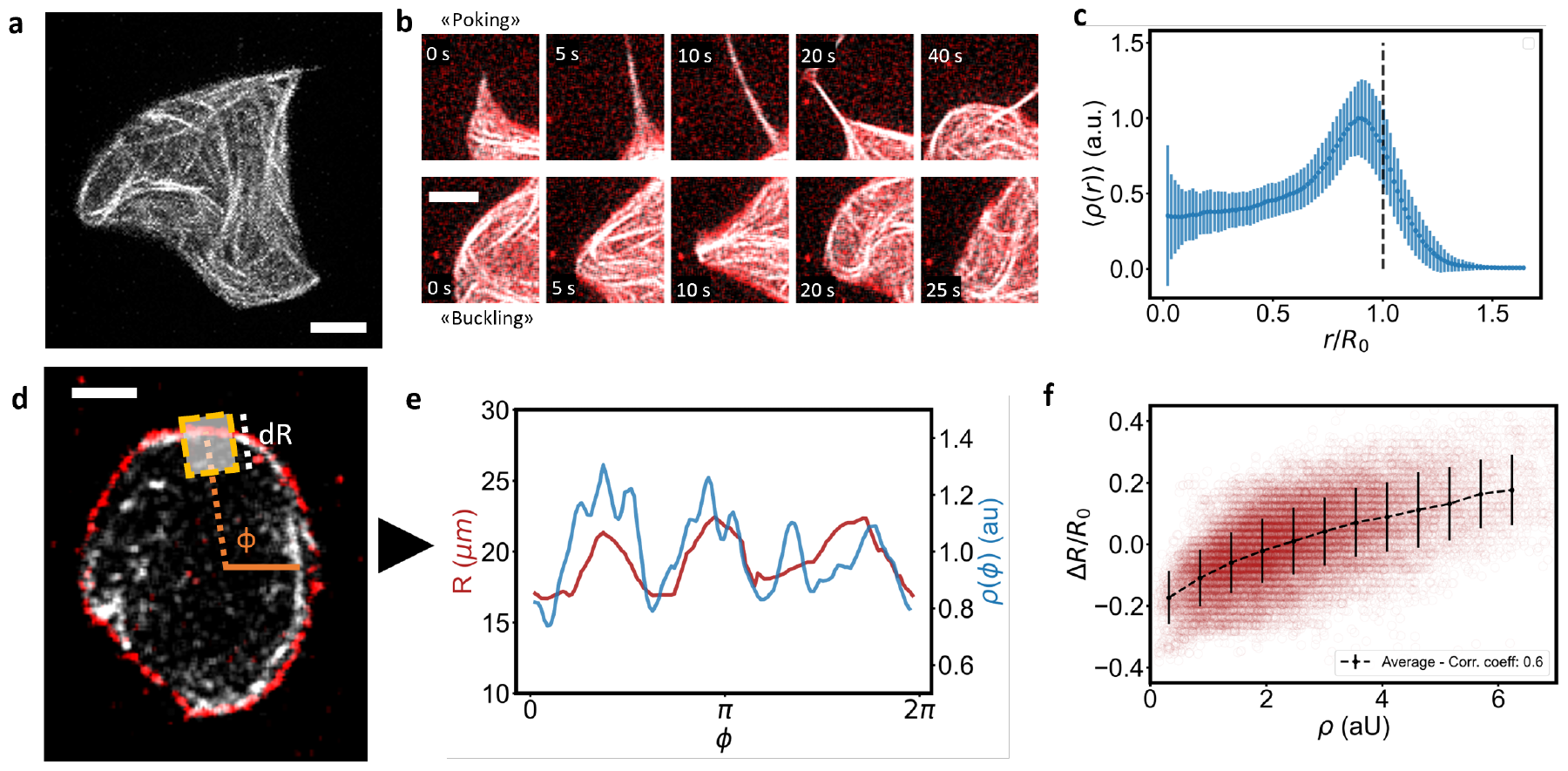
Microtubules act on the membrane to induce shape deformations: **a**, Confocal projection of the active microtubule network, showing how its structure is correlated with the GUV’s shape. Scale bar is 20 *μm*. **b**, Confocal time series, showing how MTs deform the GUV. Bundles can “poke” the membrane (top), leading to the formation of tubes or they can (bottom) “buckle” against the membrane, inducing smoother shape deformations. An arrow indicates the buckling bundles. Total duration of the series is 5 seconds. Microtubules are marked in white, membrane in red. Scale bars are 5 *μm*. **c**, Radial intensity profile of the microtubule density *ρ*(*r*), showing accumulation close to the membrane. Dashed line indicates the mean radius of the GUV over time. Errorbars indicate the standard deviation inside each bin. **d**, The microtubule intensity *ρ*(*ϕ, t*) is obtained by averaging the microtubule fluorescent intensity in a box centered at the membrane position *R*(*ϕ, t*) and with size ∆*R* = 2*μm* to obtain the angular distribution of microtubules along the membrane (not to scale in the picture). Scale bar is 20 *μm*. **e**, Plot of the microtubule density *ρ* (blue) and the local deformation ∆*R* (red) along the membrane, showing correlations between the two. A higher density leads to a higher deformation. **f**, Scatter plot of the correlation between the density and the deformations along the whole trajectory of a GUV. Black line indicates the average inside equidistant bins of the density and show a correlation coefficient of 0.6.

We identify two main ways the microtubules can push against the membrane: when they extend and push radially against the membrane, they lead to transient, tube-like deformations (“poking” behavior, Fig. 3.b top, Movie 5). Such tubes are transient and retract. When instead the bundles approach the membrane tangentially, they produce long-wavelength deformations. These latter are due to bundles buckling against the membrane (“buckling” behavior) (Fig. 3.b bottom, Movie 6). In both cases, the deformations are tightly correlated to the local activity of the MT network.

Based on these observations, we postulate that the temporal correlation of membrane deformations is connected to the dynamics of the microtubule bundles. Under the assumption that the activity of filaments is proportional to their number, irrespective of their orientation, we can then use the MT density as a proxy for the force they exert. We then extract from fluorescence movies the intensity of microtubules *ρ*(*r, ϕ, t*) where *r* is the distance from the GUV center. We find that microtubules are highly inhomogeneously distributed in space. Differently from previous systems encapsulating active microtubules [26], bundles are not confined on the surface, nor do they cover it completely. They are hence exempt from topological constraints on their alignment. Yet, while able to cross the interior, we find that filaments are still mostly concentrated along the membrane’s surface due to their extensile behavior and the radial density of MTs is peaked at *r* ≈ *R*_0_ (Fig. 3.c).

Shape changes of the membrane are hence driven by the local organization and activity of the microtubules, that directly push against the membrane and deform it. Thus, we can capture the relevant dynamics of filaments and their interplay with the membrane by reducing our observable to an angular density of microtubules *ρ*(*ϕ, t*) := *ρ*(*r* ≈ *R*_0_, *ϕ, t*) computed only in the vicinity of the membrane (Fig. 3.d). From it, directly correlating membrane deformations with the local MT density, we confirm that filaments are also highly concentrated where membrane deformations are larger than average (Fig. 3.e-f). We then turn to the dynamics of the MTs. By tracking the flow of microtubules along the membrane (Fig. S6.A-B), we observed that clusters of highly-concentrated microtubules travel along the membrane with a speed of *v* ≈ 1 *μm/s* (Fig. 4.a, Movie 7-8), transporting with them the membrane deformations they induce (Fig 4.b). This gives rise to transient deformation waves that travel, merge or split (Fig. 4.c, Movie 4), and then switch direction or dissolve after a typical time. These transient waves are due to the tendency of the microtubule bundles to extend, hence deforming the membrane. The resulting increase in membrane tension deflects the motion of microtubules, thus turning extensile activity into motion along the membrane. On average, 70 % of the kinetic energy is in a direction tangential to the membrane (Fig. S6.C). It follows that the dynamics of the membrane is due to microtubules deforming it due to their extensile-based pushing and then moving along the GUV’s perimeter due to confinement. In doing that, they transport these deformations along the membrane, as confirmed by the correlation between radial deformations, microtubule density along the membrane and their tangential speed which propagates the deformations (Fig 4.d(i)-d(iii)).

**Figure 4.**
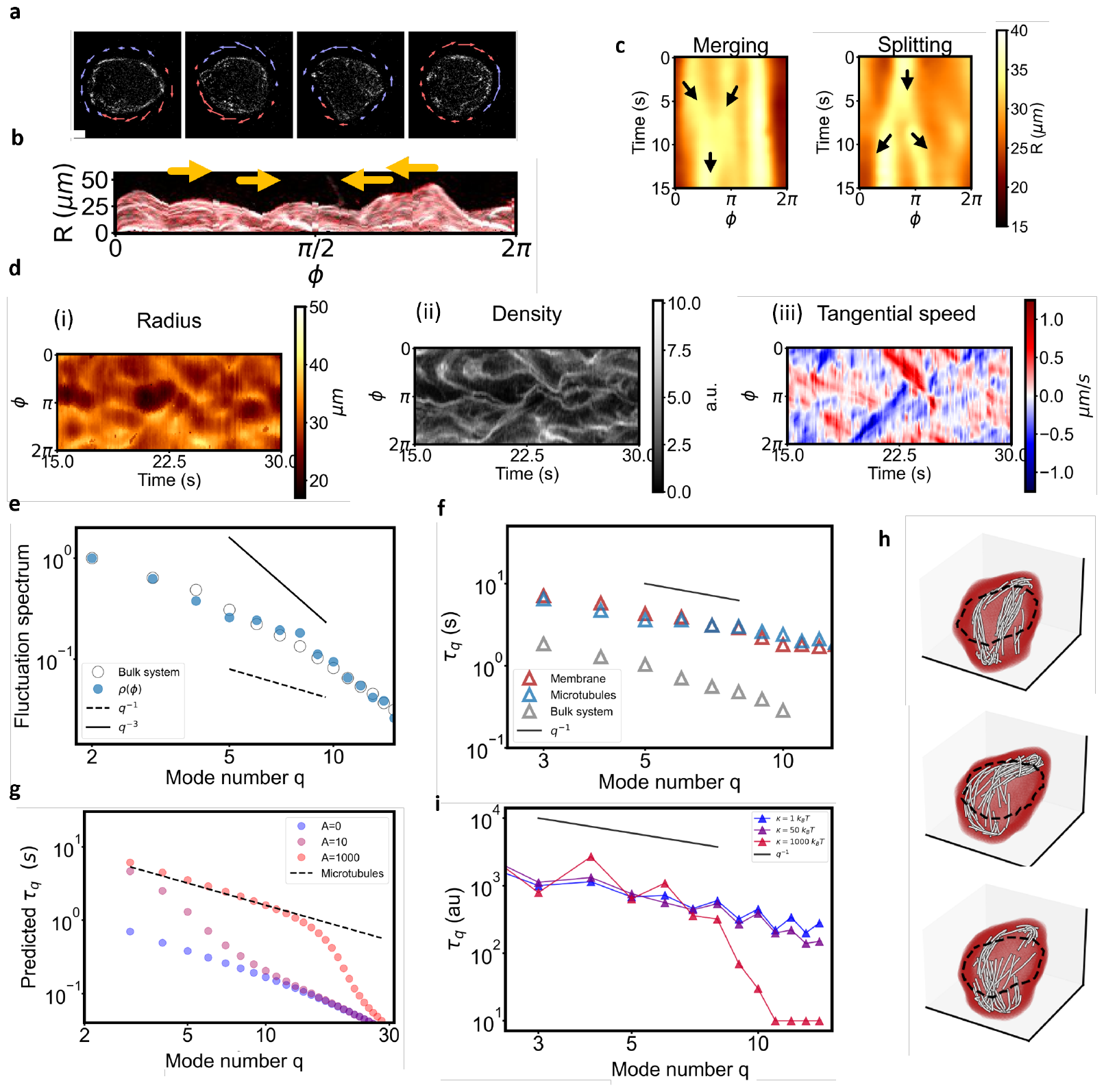
The dynamics of the microtubules sets the dynamics of membrane deformations: **a**, Microtubule flow along the membrane at four different consecutive times, showing that the flow goes both clockwise (red) and counter-clockwise (blue), and is organized in domains of similar flow (transient waves) which move, collide and rearrange over time. Scale bar is 20 *μm*. **b**, By visualizing the vesicle using confocal images projected in the (*r, ϕ*) plane, one can see microtubule-driven membrane deformations that travel along the membrane, transported by the active flow (indicated by arrows). **c**, Kymographs of the membrane deformations *R*(*ϕ, t*) showing areas of high deformations propagating in time and merging (left) or splitting (right). Arrows indicate the direction of motion. **d**, Kymograph of membrane deformations (i, left), microtubule density (ii, center) and tangential flow along the membrane (iii, right) showing high correlation and indicating how flow transports the MT density, which in turn induces the membrane deformations. **e**, Spatial fluctuations spectrum for the microtubule density *ρ*(*ϕ*) both under GUV confinement but close to the membrane (full blue circles) and in bulk (open grey circles). Both spectra are normalized so that they have a value of 1 at *q* = 2. The two quantities show a similar decay. **f**, Correlation times of deformations at each mode *q* for the membrane (red, as in 2). **g** compared to those extracted from the spectral description of the microtubule density *ρ*(*ϕ, t*) (blue). The two curves are comparable with each other and display the same scaling. The same spectral analysis, performed on the bulk system, shows a similar scaling but faster decorrelation. **g**, Solution of the Langevin equation 3 for the correlation times at three different values of the active force *F*_*lm*_. While a passive membrane (*A* = 0, blue) shows the decay predicted at equilibrium, as the force is increased (*A* = 1000, red) the membrane decay times sync to those imposed by the active force (dashed line indicates the imposed force persistence time, *T*_*lm*_ ∼ 1*/l*). The parameters used are *κ* = 10*k*_*B*_*T, R*_0_ = 25 *μm*, 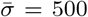, assuming the viscosity of water and with *T*_*lm*_ = 16*/l s* and *ρ*_*lm*_ ∼ *e*^−*l/*6^. **h**, Snapshots of the particle-based simulations of a GUV containing active filaments. Simulations recapitulate the results and show large membrane deformations. **i**, For stiff vesicles (*κ* = 1000*k*_*B*_ *T*), only the longest-wavelength modes show enhanced relaxation times at high activity, whereas at lower values of *κ* more modes are enhanced.

To quantitatively understand the link between microtubules and membrane fluctuations, we used again a Fourier series expansion of *ρ*(*ϕ, t*). This allowed us to compute both the density fluctuation spectrum and its temporal decay for each mode. The method is similar to what was applied to the membrane, but now focused on the active microtubule network pushing on the membrane. The fluctuation spectrum for the microtubule density, ⟨|*ρ*_*q*_|^2^⟩, as shown in Fig 4.e (blue), is distinct from that of the membrane. It aligns with what we see for an un-confined MT active gel (shown in gray, see also in Fig. S5). This suggests that the angular filament density within GUVs is arranged similarly to its distribution in bulk. This specific spatial arrangement of MTs however, when acting on the membrane, while scaling differently than equatorial fluctuations, it does influence them, resulting in the observed membrane fluctuations *q*-dependent decay. What is particularly notable is that, conversely, the microtubule density also exhibits *q*-dependent decay times 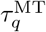 which are consistent in both magnitude and scaling with the decay times observed for the membrane fluctuations 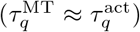. This parallelism suggests that the active microtubule fluid determines the membrane’s fluctuation timescale (see Fig. 4.f and Fig. S4.B). This elucidates the observed separation between the membrane’s spatial and temporal correlations, as the latter are now solely set by the active MT network.

Moreover, again examining a similar microtubule system in bulk under identical conditions, we found a similar scaling of the correlation times but with faster decay times (around ten times lower than inside GUVs). This indicates that the confinement induced by the GUV slows down the system’s dynamics (Fig. 4.f, gray). Hence soft confinement, while not modifying its angular distribution, extends the microtubule density’s correlation timescale. We attribute this effect to the membrane acting as barrier which redirects any radial flow of the network tangentially, thus prolonging its persistence in time.

## A Phenomenological model recapitulates the observed dynamics

The observed dynamics is qualitatively captured by a Langevin equation, describing the evolution of the GUV’s surface, where the Fourier modes *u*_*q*_ are extended to coefficients *f*_*lm*_, describing spherical modes (*l, m*) [13], under the influence of an active force *F*_*lm*_(*t*) correlated in time. The equation is given by

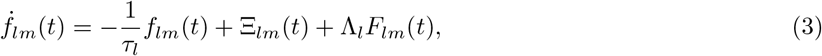

where *τ*_*l*_ is a passive response time, only dependent on *l*, Ξ_*lm*_ is a thermal noise and Λ_*l*_ is the Oseen tensor (see SI). Equation 3 can be easily understood in terms of each mode being excited by the active force *F*_*lm*_ (which we further assume proportional to the microtubule density through a constant *A*, that also serves as an activity parameter) but responding to it through the passive term −(1*/τ*_*l*_)*f*_*lm*_. The spectral characteristics of the active force are 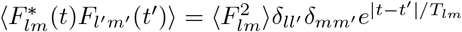, with *T*_*lm*_ indicating the persistence time of the force.

This description naturally emerges from a microscopic model in which filaments push persistently on the GUV’s surface, and its details are validated by the experimentally measured correlation functions of the microtubule density (Fig. S4). We impose, as observed in experiments, *T*_*lm*_ ∼ 1*/l*.

By fully solving the model, the final fluctuation spectrum is found to be a sum of a passive component with an active one [21], which dominates at high enough forces. Weaker activity instead leads to small modifications in the membrane decay [14, 48]. When *A* = 0 we recover the expected passive fluctuations spectrum. For *A >* 0, we also recover that the scaling of membrane fluctuations is different from that of its correlation times. The latter, as expected, synchronize with the timescales of the active force when activity is increased (Fig. 4.g). Hence the presence of a correlated active term acting on the membrane explains the observed spatio-temporal symmetry breaking. While the synchronization of timescales is a general result as long as activity is high enough and *T*_*lm*_ ≫ *τ*_*l*_, the scaling of spatial fluctuations depends on several microscopic parameters, including the mechanical properties of the GUV, the correlation of the active force and its spatial organization, the latter being difficult to extract from experimental data. Using reasonable assumptions, motivated by experimental results, we recover the behavior of the observed fluctuation spectra for active GUVs (Fig. S7).

In addition, we performed particle-based simulations of active filaments inside a deformable membrane (Fig. 4.h), for a range of vesicle bending modulus values. We find that filaments accumulate at the vesicle boundary, resulting in interaction forces that are highly correlated with filament density. Force fluctuations in different spherical harmonic modes (*l, m*) are approximately uncorrelated and exponentially decaying in time, thus further validating our assumptions. Also in this case, as predicted, the final result is that vesicle height fluctuations are enhanced by active forces, with vesicle relaxation times following those of the interaction forces (Figs. S8-S9-S10). Moreover, the simulations allow us to probe the role of vesicle deformability: for stiff vesicles, only the longest-wavelength modes show enhanced height fluctuations and relaxation times as activity is increased (Fig. 4.i), underlining the key role of vesicle rigidity in modulating the magnitude and the lifetime of active deformations.

The observed fluctuations in the membrane and microtubule dynamics hence reflect a feedback in which soft confinement modifies the spatio-temporal microtubule organization, which in turn dictates large membrane fluctuations around the equilibrium spherical shape. The resulting activity of the MT network acting on the membrane is correlated in time and drives the dynamics of membrane fluctuations accordingly.

We have shown that an active cytoskeleton, which spans the volume of a vesicle and pushes on its membrane, leads to seemingly irregular large shape transformations. Yet, the vesicle deformations still exhibit well-defined decays in their power spectra, arising from an intricate coupling between active transport processes within the cytoskeleton and soft confinement by GUVs. Specifically, once the strength and temporal organization of the active system dominate over the passive mechanical properties of the membrane, the typical timescale of the active system sets the membrane fluctuation timescale. Such a direct link between the membrane and the underlying active system implies that the temporal dynamics of the membrane fluctuations, rather than its spatial counterpart, carries with it the signature of the underlying active system [15, 16, 18, 19]. Our results, extending the equilibrium theory of membrane fluctuations [11–13] to active systems, provide a benchmark to study the interplay between a membrane and an actively reorganizing cytoskeleton in more complex model systems and for living cells. Understanding this dynamics is a also fundamental step in the programs of reconstituting a synthetic cell and developing biologically-inspired soft-robots. The understanding of active membrane fluctuations paves the way to the design of systems whose activity might be harnessed to obtain shape-morphing and self-propelled synthetic cells capable of exploring complex environments.

## I. ACKNOWLEDGEMENTS

AS and ARB acknowledge the support by the European Research Council (ERC) under the European Union’s Horizon 2020 research and innovation programme (grant agreement no. 810104-PoInt). This research was supported in part by the National Science Foundation under Grant Nos. NSF PHY-1748958, NSF DMR-1855914 (SU and MFH), NSF DMR-2202353 (AB) and the Brandeis Center for Bioinspired Soft Materials, an NSF MRSEC, DMR-2011846 (LBF, MFH, AB). We also acknowledge computational support from NSF XSEDE computing resources allocation TG-MCB090163 (Expanse, Anvil, and Bridges-2) and the Brandeis HPCC which is partially supported by the NSF through DMR-MRSEC 2011846 and OAC-1920147. AS and ARB acknowledge the Brandeis MRSEC for shipping microtubules and kinesin (Grant No. MRSEC-DMR-2011846). The authors thank Philip Bleicher for help with the purification of anillin.

## II. AUTHOR CONTRIBUTIONS

AS designed and performed experiments, analyzed the data and wrote the manuscript. HF designed research, analyzed the data and wrote the manuscript. SU, LF and MSE performed and analysed simulations. PV, AB, MH designed research. ARB designed research and wrote the manuscript. All authors revised the manuscript. AS and HF contributed equally to this work.

## III. METHODS

### A. Buffers and proteins

M2B is 80 mM Pipes (pH 6.8), 2mM *MgCl*_2_ and 1 mM EGTA. 3.2 mM of *MgCl*_2_ is added at the end from a 67 mM stock already diluted in M2B.

Lipids (TexasRed-DHPE and DOPC, Avanti Polar Lipids) are bought from ThermoFisher. Anillin is purified in the laboratory from its sequence (see SI). Silicon and mineral oil are purchased from Sigma.

### B. Encapsulation using cDICE

Vesicles are produced using the cDICE method[42, 49] consisting briefly of letting droplets of the active mixture cross a layer of oil (silicon and mineral oil, from Sigma) containing lipids, in order to coat them with a membrane. The droplets is produced inserting a capillary in a 3D-printed rotating chamber[27]. Finally, the vesicles accumulate in a buffered aqueous solution, osmotically matched to the active mix using glucose. The preparation of the mixture is as follows: the desired concentration of stabilized microtubules is mixed in M2B together with the desired concentration of kinesins, ANLN and ATP. The mixture also contains a scavenging system (10 U/mL glucose-oxidase, 1 kU/mL catalase, from Sigma), glucose (3 mg/ml), an ATP regeneration system (18.2 units/mL of creatine-phosphokinase and 9 mM of creatine-phosphate, from Sigma) and 5 mM DTT. It is made sure that the final concentration of salts is exactly the one expected for M2B by correcting using a 10x M2B preparation. Both the stock of MTs and the mixture are kept at room temperature to avoid depolymerization. After mixing, we wait for 5 minutes for the active fluid to assemble. Its osmotic pressure is measured with a Gondotec osmometer and a glucose solution with comparable osmotic pressure is also prepared. Finally, the cDICE encapsulation is carried out at room temperature. We use a capillary with a diameter of 40 *μm* to allow for big vesicles and fast encapsulation (5 minutes). The vesicles are then harvested, transferred to a glass coverslip coated with 1 mg/ml Bovine serum albumine (Sigma) and observed at a microscope.

### C. Imaging

Vesicles are acquired using a confocal microscope (Leica DMi5, 63x objective NA 1.4) equipped with a resonant scanner. Long time series of the equatorial plane are acquired using a 256×256 image size with a time-interval of 35 to 250 ms (around 2000 frames). The pinhole was set to 1 Airy unit. Both the membrane and the microtubule channel are acquired at the same time. Three dimensional stacks are also acquired at different resolution and time interval.

### D. Extraction of GUV’s contour and microtubule density

The images of the membrane channel are thresholded and the GUVs’ contour is obtained using custom written Python3 code. Briefly, using as a reference the center of the GUV, the space is divided in overlapping angular segments of amplitude *da* = 0.25 rad and separate by *dϕ* = 0.15, inside which the radial position of the point with maximum intensity of the membrane is found, thus obtaining a discretization of *R*(*ϕ*). The same logic is applied to the microtubule intensity to obtain *ρ*(*ϕ, r*), which can then be averaged inside a box of width *da* and contained between *R*(*ϕ*) and *R*(*ϕ*) − *dR* with *dR* = 2*μm* to obtain *ρ*(*ϕ*) in proximity of the membrane.

### E. Kurtosis analysis

The extracted contours are splitted in series of length *τ*_avg_ and from each series the histogram of the deformations (*R* − *R*_0_)*/R*_0_ (where *R*_0_ is the mean radius of the GUV over time) is extracted. The kurtosis for each of the resulting distribution is then computed and normalized so that the expected value for a Gaussian is 0 (excess kurtosis). For each value of *τ*_avg_ we then obtain a list of kurtosis which are collected and from which their average, standard deviation and minimum and maximum value are computed.

### F. Fluctuation spectra and decay times

The discretized version of *R*(*ϕ*) and *ρ*(*ϕ*) are, at each time point, expanded in Fourier series to obtain the complex coefficients *u*_*q*_(*t*). To do that, we compute the integrals

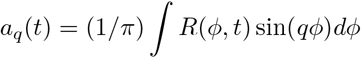

and

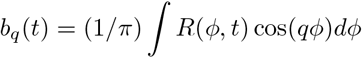

using the trapezoidal rule.

From it, we obtain *u*_*q*_ = (−*a*_*q*_, *b*_*q*_).

The variance of *u*_*q*_(*t*) yields the spatial spectrum whereas it correlation function is used to extract decay times. The decay time is defined as the time at which the correlation as decreased below 1*/e* of the initial value to compare decays which are not strictly speaking expoentials.

### G. Broken Detailed Balance

We extract the microscopic configurations of GUVs which here to the shapes defined by different Fourier modes, discretized. The probability of a given configuration is defined as the ratio of the time spent at a given shape configuration. The currents across box boundaries determined by counting statistics that is the transitions between boxes yields the the probability current **j**. A nonzero value of its contour integral 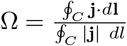, indicates a system out of equilibrium where *C* is a cycle. The flux is normalized hence dimensionless. More details about the method can be found in [46, 47]. The z-score of each flux is obtained by collecting all values of Ω across different cycles, taking their average and checking how many standard deviations away from Ω = 0 it is.

### H. Analysis of MT flow

To extract the flow of microtubules inside GUVs the movies of the MT channel at the equatorial plane are analysed with an optical flow algorithm using a custom Python3 script. Roughly, the intensity is followed over time extracting its flow, then the flow close to the membrane (using the procedure detailed above for the MT density) is averaged over 1 *s* and decomposed into tangential and radial components by a scalar product with a unit vector starting from the center of the GUV and extending radially and its normal counterpart (tangential).

#### I. Bulk experiments

To perform bulk experiments, the same mixture is injected inside a 10 *μl* microscopy chamber composed of a glass slide and a coverslip separated by a layer of parafilm. The slides and coverslips are passivated with polyacrylamide [41]. The Fourier analysis of the bulk fluid is detailed in SI.

## Movie Legends

**Movie 1:** Confocal projection of a vesicle containing an active MT network and undergoing shape changes. The vesicle contains 0.8 mg/ml microtubules, 120 nM Kinesin and 1.5 μM anillin. Microtubules in white, membrane in red. Time interval is 2.5 s, scale bar is 50 μm.

**Movie 2:** Confocal three-dimensional reconstruction of a vesicle containing an active MT network. The vesicle contains 0.8 mg/ml microtubules, 120 nM Kinesin and 1.5 μM anillin. Microtubules in white, membrane in red. Time interval is 2.5 s, scale bar is 50 μm.

**Movie 3:** Equatorial fluctuations of an active vesicle. Microtubules in white, membrane in red. Time interval is 100 ms, scale bar is 50 μm.

**Movie 4:** Confocal three-dimensional reconstruction of the active MT network inside an active vesicle. Left pane shows the Z-projection over time, whereas the two other panes show a three-dimensional reconstruction from two different points of view. The vesicle contains 0.8 mg/ml microtubules, 120 nM Kinesin and 1.5 μM anillin. Time interval is 2.5 s, scale bar is 50 μm.

**Movie 5**: Detail of a vesicle undergoing “poking”. The vesicle contains 0.8 mg/ml microtubules, 120 nM Kinesin and 1.5 μM anillin. Membrane in red, microtubules in white. Time interval is 2 s.

**Movie 6:** Detail of a vesicle undergoing “buckling”. The vesicle contains 0.8 mg/ml microtubules, 120 nM Kinesin and 1.5 μM anillin. Membrane in red, microtubules in white. Time interval is 2 s.

**Movie 7:** Flow of microtubules in the proximity of the membrane inside an active vesicle. Blue arrows indicate counter-clockwise flow, blue arrows indicate clockwise flow. Microtubules are in white. Time interval is 4 s, scale bar is 20 μm.

**Movie 8:** A fluctuating vesicle is projected in the (R, ϕ) plane to highlight the motion of membrane fluctuations along the membrane. Time interval is 2.5 s.

