## Supplementary Information for "Active membrane deformations of a minimal synthetic cell"

### Supporting Information

November 14, 2023

### 1 Methods and Material

#### 1.1 Anillin purification

Anillin (ANLN) is an actin-binding protein that bundles filaments. We use it as an (unspecific) microtubule crosslinker. We clone ANLN using the following (His-Tag containing) amino acid sequence:

MGSSHHHHHHSSGLVPRGSHMDPFTEKLLERTRARRENLQKKMADRPTAGTRTAAL
NKRPREPLLEANHQPPAPAEEAKPSSKPSPSKRRCSDNASTPDAGAENKQPKTPEL
PKTELSAVASHQQLRATNQTPQVSLLSSDKELTASDVKDASSVKTRMQKLADQRRY
WDNNVSPSSSPPAHVPPKDIIVSPPKPQIPDVGNDTPVGRRGRFANLAATIGSWED
DLSHPFVKPNNKQEKPGTACLSKESTTSSASASMNSRSVKQDTTSCSQRPKDTTVN
KAVCSGQLKNILPASKPASSVASTEVSGKSKPLAIKSPTVVTSKPNENVLPASSSL
KPVSANSSPQKTERPASRIYSYQSASARNELNNNTPVQTQQKDKVATSGGVGIKSF
LERFGEKCQEHSPAPLNQGHRTAVLTPNTKSIQERLLKQNDISSTALEHHHHHH

We use a pET28b(+) vector. Plasmids are amplified with chemically competent E. coli XL-1 blue cells for DNA production and E. coli BL-21 CodonPlus for protein expression, both purchased from Agilent Technologies.

### 2 Experimental methods

### 3 Fluctuation spectroscopy and data analysis

#### 3.1 Extraction of GUV's contour

We acquire movies of the equatorial plane of fluctuating vesicles, starting from the membrane channel. The procedure is the same for active or passive vesicles. To extract the contour, we use previously developed algorithms [3]. The program was coded and compiled on MATLAB (Mathworks, USA) with the help of inbuilt MATLAB functions or using Python3 scripts. The software performs

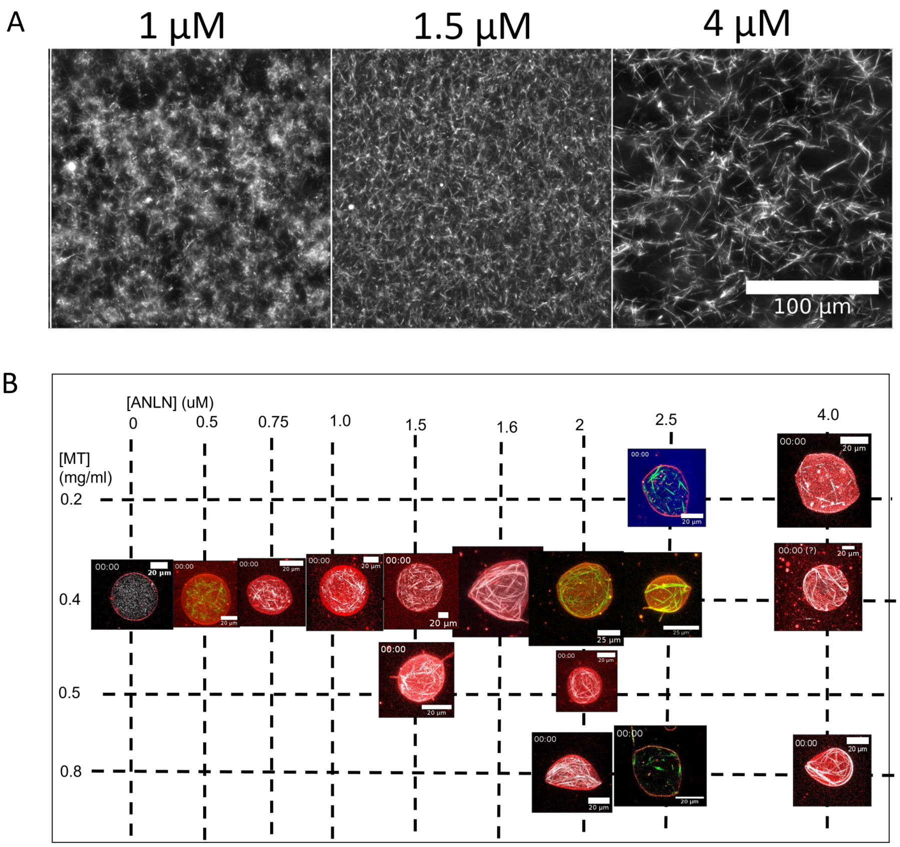


Supp.Fig. 1: A)Anillin bundles microtubules. Adding different concentrations of anillin to a mixture of 8 mg/ml of microtubules leads to the formation of bundles. B) Active vesicles are observed for a range of ANLN and MT concentration. All GUVs contain 60 nM Kinesin in addition to the indicated concentrations.

three important steps to detect the contour; (i) image processing, (ii) pixel and sub-pixel resolution contour detection and (iii) fitting the vesicle contour by a Fourier series. The details are as follows:

After acquiring the raw confocal fluorescence images of the membrane, we implement an inbuilt MATLAB sobel disk filter $fspecial\ ('sobel')$ and image normalization to increase the contrast of the contour. Initially, an estimated radius is determined by converting 10-100 consecutive contours into binary images. A skeletonization ($bwmorph$) and circle fitting algorithm (least square method) is used to determine an approximate radius of the vesicle. The first approximated radius is used to implement an inbuilt MATLAB function $imfindcircles()$. The function uses circular Hough transform to give an improved approximated radius $R_{ves}$ and first guess of the centroid of the circle. This forms our preliminary image detection technique. The analysis becomes more precise by using the approximated centroid and $R_{ves}$. The cartesian plane in the MATLAB grid is converted to polar grid where the centroid is placed at the origin. The software proceeds to find the pixel location of minimum intensity in N regions of size $d\phi$=$2\pi/N$ (wedges). It is to be noted that N is determined in powers of 2 to implement the Discrete Fast Fourier Transform (FFT) algorithm. The new centroid is determined by averaging the coordinates of the minimum pixel location. The improved approximated radius is found by determining the mean of each found pixel location from the new centroid. This process is iterated until the centroid point converges with a difference 0.001%. It is important to note that the contour is contained within an annulus (Region of Interest) of inner radius $R_i$ and outer radius $R_o$; $R_i$<$R_{ves}$<$R_o$. This optimizes the software to work faster and prevents the algorithm from finding some defects or impurities in the images as the pixel location of the minimum intensity. After getting a converged centroid, the same algorithm of finding pixel location of minimum intensity is utilized. This is the gross signal of the contour with a pixel resolution accuracy. For the sub-pixel accuracy detection, we have implemented the algorithm from Gracia et al [4]. In summary, the sub-pixel algorithm works as follows: we perform a fitting of the gray value intensity profile around the gross location of the contour points. This yields sub-pixel accuracy of the Fourier signal. The pixel accuracy algorithm detects the minimum point of the valley of the profile. The sub-pixel resolution contour detection is determined by fitting two slope lines around the minimum intensity pixel.

##### 3.2 Fluctuation spectra and decay times

To perform flickering spectroscopy, we use a previously developed method as detailed in [3]. In summary, a time series of fluctuating vesicles at the equatorial cross section is recorded. The fluctuating contour is represented in Fourier modes, $r(\phi) = R\left(1 + \sum_q u_q(t)\exp(iq\phi)\right)$ using a Fast Fourier Transform or explicitly performing the discrete integrals

$$a_q(t) = (1/\pi)\int R(\phi,t)\sin(q\phi)d\phi$$

and

$$b_q(t) = (1/\pi)\int R(\phi,t)\cos(q\phi)d\phi$$

using the trapezoidal rule (*trapz()* from the Numpy library of Python3) from which we obtain $u_q = (-a_q, b_q)$.

The amplitude of the fluctuations $u_q$ can be presented with mean square amplitude that depends on the membrane bending rigidity $\kappa$ and the tension $\sigma$, $\langle|u_q|^2\rangle \sim \frac{k_BT}{\kappa(q^3+\bar{\sigma}q)}$, where $k_BT$ is the thermal energy ($k_B$ is the Boltzmann constant and $T$ is the temperature), and $\bar{\sigma} = \sigma R^2/\kappa$. Images are acquired with confocal microscopy at 10-30 fps for 5-10 mins. Only vesicles with low tension value ranging from $10^{-7} - 10^{-10}$ N/m are chosen. This results in a

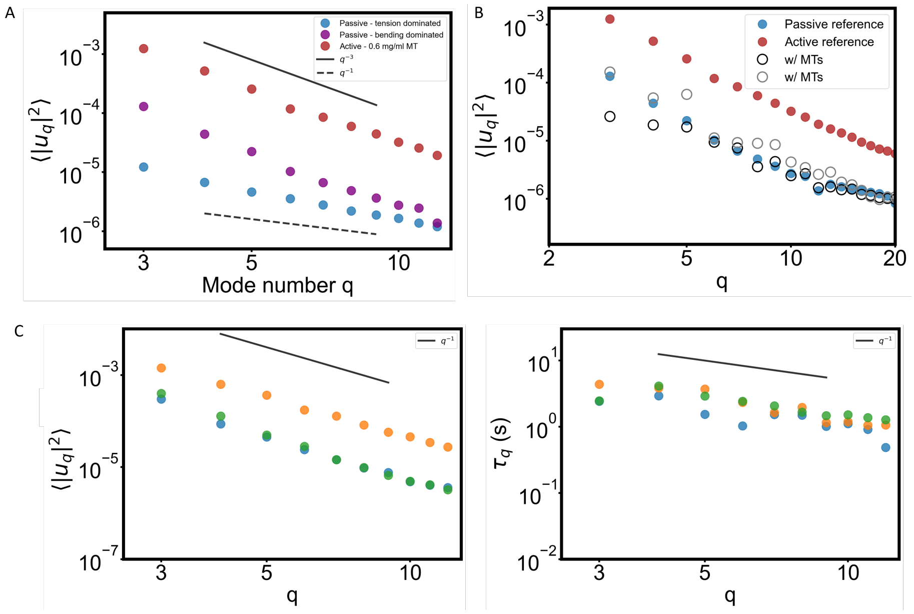


Supp.Fig. 2: A) Fluctuation spectra of an active and two passive GUVs (ref Fig. 2 of main text). The blue dots indicate a tension-dominated GUV with the spectrum scaling as $\sim 1/q$ obtained by choosing bigger GUVs so that the crossover mode between bending and tension-dominated regime is observable. B) Fluctuation spectra for two passive vesicles encapsulating microtubules but without crosslinkers and motors, showing that passive, unbundled microtubules do not significantly modify the fluctuations. A purely passive (blue)rs and an active (red) spectra are shown as references. C-D) Fluctuation spectra (C) and correlation times (D) of three different active GUVs, showing comparable scaling and timescale.

small cross over mode given by $q_c = \sqrt{\bar{\sigma}}$ where the shape fluctuation modes are dominated by bending rigidity. We have ignored the ellipsoidal mode ($q$ =2) as it is weighted with most excess area which leads to fluctuations with an increased amplitude.

Using the same time series data, one can also compute the temporal auto-correlation function which gives information about time evolution of the modes, $\langle u_q(0)u_q^*(t)\rangle = \langle |u_q|^2\rangle \exp(-t/\tau_q)$. If $q \gg 1$, the correlation time tends to that of a planar membrane $\tau_q^{-1} = \kappa\left(q^3 + \bar{\sigma}q\right)/4R_0^3\eta$. For tensionless vesicles, the correlation time would scale as $\tau_q^{-1} \sim \kappa q^3/4R_0^3\eta$.

#### 3.3 Broken Detailed Balance

The non-equilibrium nature of the vesicle fluctuations can be quantified by the method of broken detailed balance. The method states that for a system in equilibrium, driven by thermal forces, observable microscopic configurations must be pairwise balanced. This refers to equal likelihood for the forward and backward transition between two processes. In our case, the microscopic configurations refer to the shapes defined by different Fourier modes. A non-equilibrium system, however, would display a probability flux cycle in the phase space of shapes; this refers to unequal rate of transitions between forward and backward shape changes. On the other hand, the probability is defined as the fraction of the time spent at a given shape configuration. A nonzero value of the contour integral of the probability current, $\Omega = \frac{\oint_C \mathbf{j}\cdot d\mathbf{l}}{\oint_C |\mathbf{j}|\ dl}$, indicates a system out of equilibrium. More details about the method can be found in [2, 5].

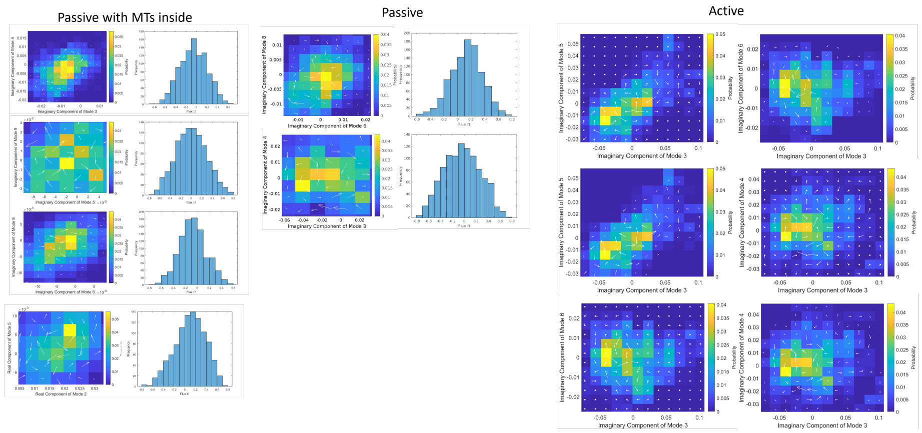


Supp.Fig. 3: Map and histogram of the flux $\Omega$ between different modes for two passive vesicles, one containing MTs but no motors (left), on empty (center) and for an active (right) one. Only active ones show overall net flux between modes. The colormap demonstrates the probability density for the two chosen Fourier modes, the size of the arrows indicate the currents across box boundaries determined by counting statistics – that is, the transitions between boxes.

#### 3.4 Analysis of the angular density of microtubules

From the position of the membrane $R(\phi, t)$ we compute the angular density of microtubules close to the membrane $\rho(\phi, t)$ by averaging the microtubule fluorescence intensity inside a box of size 2 $\mu m$ centered at $R(\phi, t)$. The density is normalized by the mean intensity over all boxes. The spacing between boxes is the same as the angular resolution $d\phi$ used to compute the membrane position.

#### 3.5 Analysis of bulk fluid

To analyse the bulk fluid and compare it to the GUV, starting from time-lapse movies of the bulk fluid the angular intensity of microtubules is computed by

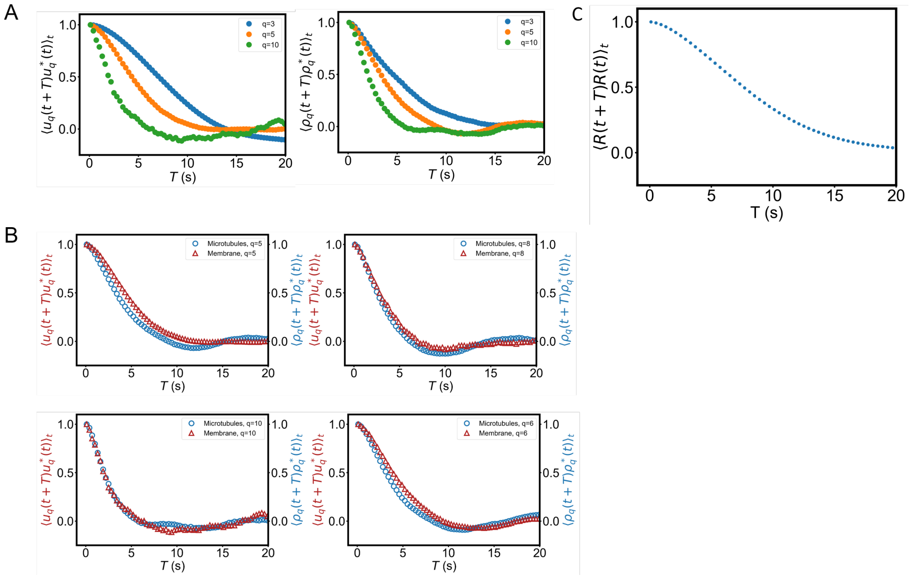


Supp.Fig. 4: A) Correlation function over time for modes $q = 3,\ 5,\ 10$ of both the membrane coefficients $u_q$ (left) and the microtubules density $\rho_q$ (right) showing they decay similarly and they have a roughly exponential decay especially at higher modes. B) Comparison for different values of $q$ between the membrane (red) and the density (blue) correlation functions, showing that in the presence of activity both quantities decay in the same way. C) Correlation function in real space of the radial deformations, showing the radial deformations are correlated for $\approx 5$ seconds in real space.

measuring the fluorescence intensity inside virtual circles of radius $\approx 25\ \mu m$ and thickness $dR = 2\ \mu m$. The virtual circles mimic the procedure based on the GUV membrane as above. Having acquired the angular density $\rho^B(\phi, t)$, it is decomposed in Fourier modes as above and the spectrum and correlation times are computed (Fig. S5).

### 4 Theoretical model and simulations

#### 4.1 Langevin equation

Take a GUV with bending rigidity $\kappa$, radius $R_0$ and tension $\sigma$ and hence with a reduced tension $\bar{\sigma} = \sigma R_0^2/\kappa$ We start from a description of the membrane in 3D as a sum of spherical harmonics (denoted as $Y_{lm}(\phi, \theta)$) given by (following [3]):

$$R(\phi, \theta, t) = R_0 \left( 1 + \sum_{l=0}^{l_{\max}} \sum_{m=-l}^{l} f_{lm}(t)\, Y_{lm}(\phi, \theta) \right) \tag{1}$$

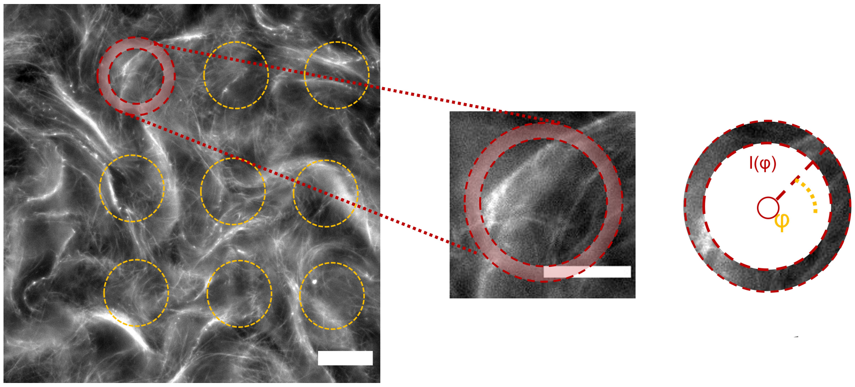


Supp.Fig. 5: Scheme of the analysis of bulk MT fluid in Fourier space. Starting from confocal planes of the bulk fluid (left), we extract the MT intensity inside circles of radius 20 $\mu m$ and thickness 2 $\mu m$ (center, right) and use it to compute the angular intensity as done for GUVs. This density is then converted in Fourier modes whose fluctuations are analysed.

where $f_{lm}(t)$ are the coefficients of the decomposition. We can set $f_{1m} = 0$ for $m = -1, 0, 1$ as this accounts for translations and $f_{00}$ is set by volume conservation and we ignore it in the following.

Using this description, the equatorial coefficients $u_q$ are obtained by summing the spherical harmonics coefficients and setting $\theta = \pi/2$ obtaining

$$u_q(t) = \sum_{l=q}^{l_{\max}} f_{lq}(t)\, Y_{lm}(\phi = 0, \theta = \pi/2) e^{-iq\phi} \tag{2}$$

(where explicit terms in $\phi$ in the definition of the spherical harmonics cancel which is equivalent to set $\phi = 0$). From this it follows that

$$\langle |u_q|^2 \rangle = \sum_{l=q}^{l_{\max}} |f_{lq}|^2\, Y_{lm}(\phi, \theta = \pi/2)\, \bar{Y_{lm}}(\phi, \theta = \pi/2) \tag{3}$$

where $\bar{Y_{lm}}$ is the complex conjugate of $Y_{lm}$.

We then assume each mode is independent and that it evolves according to the Langevin equation

$$\dot{f}_{lm}(t) = -\omega_l f_{lm}(t) + \Xi_{lm}(t) + \Lambda_l F_{lm}(t). \tag{4}$$

with:

$\omega_l$ being the inverse of a passive correlation time, given by, under the assumption that the viscosity of the GUV is the same inside and outside

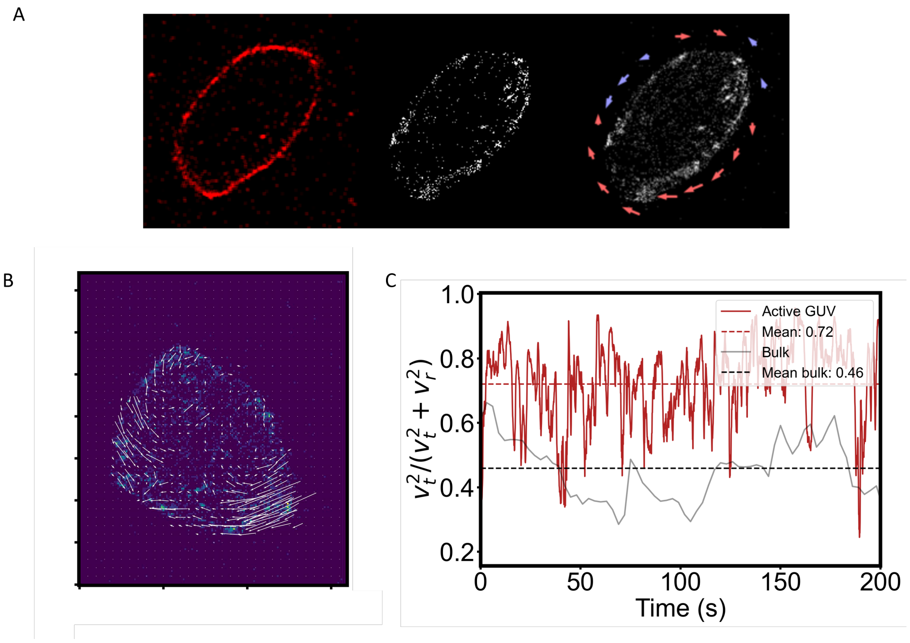


Supp.Fig. 6: A) Membrane, microtubules and extracted flow. In all cases, the membrane is identified and density and flow are computed everywhere, but then only points close to the membrane are considered. B) Example of extracted flow. C) By extracting the MT flow $\mathbf{v} = (v_x, v_y)$ inside GUVs, we can compute the ratio of the kinetic energy between the tangential speed $v_t = \mathbf{v} \cdot \mathbf{n}$ and the radial speed $v_r = \mathbf{v} \cdot \mathbf{r}$ where $\mathbf{r}$ and $\mathbf{n}$ are the radial and tangential versors ($\mathbf{n} \cdot \mathbf{r} = 0$) (shown in red for a GUV). We find that roughly ≈70% of the kinetic energy is stored in motion tangential to the membrane. In comparison, the bulk system (gray) shows as expected ≈50% (where the origin of the radial and tangential versor have been chosen at the center of the image).

$$\omega_l = \frac{\kappa(4l^3 + 6l^2 - 1)(l+2)(l-1)(l^2 + l + \bar{\sigma})}{(l^2 + l)\eta R_0^3}; \tag{5}$$

$\Xi_{lm}$ being a thermal noise with vanishing mean, delta correlated in time and and variance given by

$$\langle \Xi_{lm}^2 \rangle = (-1)^m \frac{2k_B T(l^2 + l)}{\eta R_0^3 (4l^3 + 6l^2 - 1)} \tag{6}$$

whereas all the mixed terms $(l, l')$ and $(m, m')$ vanish;

$F_{lm}$ being an unknown active force which is coupled to the membrane via the Oseen tensor $\Lambda_l$.

We note that if $F_{lm} = 0$ we recover the equations for a passive GUV [3].

We make the further assumptions that the force is proportional to the microtubule density $\rho_{lm}$ and it is exponentially correlated in time, so that

$$F_{lm} = A\rho_{lm}$$

and

$$\langle F_{lm}(t)F_{lm}(t')\rangle = \langle |F_{lm}|^2\rangle e^{-\Omega_{lm}|t-t'|} = A^2\langle|\rho_{lm}|^2\rangle e^{-\Omega_{lm}|t-t'|},$$

$\Omega_{lm}$ being the inverse of the active correlation time and $A$ is just a constant relating density and activity.

At the moment we make no further assumption and proceed to solve the Langevin equation which under the above assumptions results in

$$f_{lm}(t) = f_{lm}(0)e^{-\omega_l t} + \int_0^t e^{\omega_l(s-t)}ds\ \Xi(s)_{lm} + \int_0^t ds\ e^{\omega_l(s-t)}\Lambda_l F_{lm}(s) \quad (7)$$

As we are only interested in the steady-state values of $\langle|f_{lm}|^2\rangle$, we can take the limit

$$|f_{lm}|^2 = \lim_{t\to\infty}\int_0^t ds \int_0^t dw\ e^{s+w-2t}\ \big(\ \langle\Xi_{lm}(s)\Xi_{lm}(w)\rangle + \Lambda_l^2\langle F_{lm}(s)F_{lm}(w)\rangle\ \big) \quad (8)$$

which is clearly divided in a passive part and an active one, containing the active force. Using the properties of the noise and of the active force, the passive part yields

$$|f_{lm}|^2_{\text{passive}} = \frac{k_BT}{\kappa}\frac{1}{(l+2)(l-1)(l^2+l+\sigma_n)}$$

whereas the active part yields

$$|f_{lm}|^2_{\text{active}} = \Lambda_l^2 A^2\langle\rho_{lm}^2\rangle\frac{1}{\omega_l(\omega_l+\Omega_l)}$$

so that the equatorial plane (summing according to Eq. 3) is expected to have as a spectrum

$$\langle u_q u_q^*\rangle = \sum_{l=q}^{l_{\max}}\left(|f_{lq}|^2_{\text{passive}} + |f_{lq}|^2_{\text{active}}\right)\ |Y_{lq}(\theta=\pi/2,\phi=0)|^2$$

Similarly, we can obtain an expression for the correlation function for the equatorial modes

$$\langle u_q(t+\tau)u_q^*(t)\rangle =$$

$$\sum_{l=q}^{l_{\max}}\left(\frac{k_BT}{\kappa}\frac{1}{(l+2)(l-1)(l^2+l+\sigma_n)}e^{-\omega_l\tau} + \Lambda_l^2A^2\langle\rho_{lm}^2\rangle\frac{\Omega_q e^{-\omega_l\tau} - \omega_l e^{-\Omega_l\tau}}{\omega_l(\omega_l+\Omega_l)(\omega_l-\Omega_l)}\right)\ |Y_{lq}(\theta=\pi/2,\phi=0)|^2$$

that is a sum of exponentials, some coming from the passive and some from the active part of the temporal response.

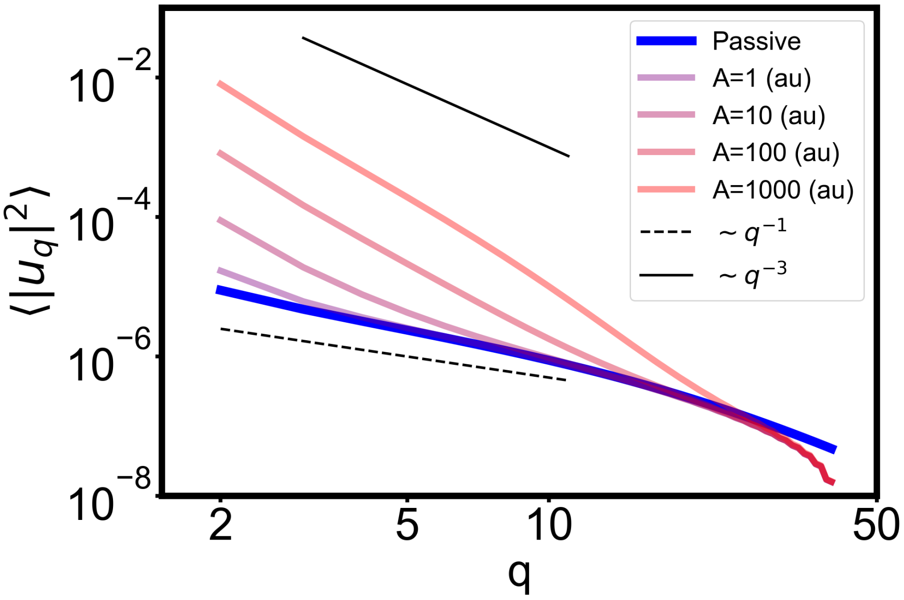


Supp.Fig. 7: Predicted fluctuation spectra at different values of the activity A for $\kappa = 15\ k_B T$.

#### 4.2 Simulations

##### 4.2.1 Overview

The fluctuations of a passive vesicle are governed by equilibrium statistical mechanics. At length scales large compared to the size of the constituent amphiphilic molecules, these fluctuations are determined by solely by the temperature and a small number of material properties (bending modulus $\kappa$ and surface tension $\sigma$.)[7, 13, 12, 8] Moreover, the fluctuations scale in a straightforward way with the wavelength at which they are probed. Concretely, if we assume that deformations $R(\theta, \phi)$ (parametrized by a polar angle $\theta$ and an azimuthal angle $\phi$) of the vesicle with respect to a perfect sphere of radius $R_0$ are small, then it is reasonable to write these deformations in a basis of spherical harmonics $Y_{lm}(\theta, \phi)$:

$$R(\theta, \phi) = R_0 \left( 1 + \sum_{l=2}^{l_{\max}} \sum_{m=-l}^{l} f_{lm} Y_{lm}(\theta, \phi) \right), \tag{9}$$

where the $l = 0$ term has been pulled out of the sum and the $l = 1$ terms, which correspond to center-of-mass translations,[12] have been excluded. The upper limit $l_{\max}$ of the first sum is set by the area of a constituent molecule compared to the vesicle area. By the fluctuation-dissipation theorem,[8] fluctuations of the spherical harmonic components $f_{lm}$ are simply proportional to their relaxation times:

$$\langle |f_{lm}|^2 \rangle \propto \tau_l. \tag{10}$$

For vesicles enclosing active filaments, this equilibrium picture breaks down. The experiments show that the mode relaxation times of such vesicles are no longer governed by their material properties as suggested by the fluctuation-dissipation theorem, but instead appear to be closely related to timescales associated with the filament organisation, as measured by filament density correlations. We performed molecular dynamics simulations of a simplified model of the experimental system to gain additional microscopic insight into the effect of active filament dynamics on membrane fluctuations.

##### 4.2.2 Model and Simulation Details

Our model for vesicles containing active filaments is very similar to a model we previously devised in Ref. [10]. We represent a vesicle as a triangulated mesh whose initial spherical configuration is generated using the polyHédronisme software package[6]. The radius of the initial, perfectly spherical vesicle is $R_0 \approx 25\sigma$, with $\sigma$ our basic unit of length. The vesicle's $N_{\rm ves} = 2432$ vertices each have diameter $\sigma_{\rm ves} \approx 1.934\sigma$. The connectivity of this mesh is fixed in our simulations.

Vertices (henceforth vesicle "beads" or "monomers") are connected to their nearest neighbors by nonlinear springs ("tethers") with potential energy:[9]

$$U_{\rm tether}(r) = \begin{cases} \frac{b\exp\left(1/(l_{c0}-r)\right)}{l_{\rm max}-r}, & r > l_{c0} \\ \frac{b\exp\left(1/(r-l_{c1})\right)}{r-l_{\rm min}}, & r < l_{c1} \\ 0, & l_{c1} \leq r \leq l_{c0}. \end{cases} \tag{11}$$

Here, $r$ is the distance between beads, $l_{\rm min} = 1.33\sigma$ and $l_{\rm max} = 2.67\sigma$ respectively set the minimum and maximum allowed distance between bonded beads, $l_{c1} = 1.66\sigma$ and $l_{c0} = 2.34\sigma$ respectively set the lower and upper limits of a "flat" region where the potential is zero, and $b = 80k_{\rm B}T\sigma$ sets the strength of the tether interaction keeping the beads bonded.

Pairs of triangles (with normal vectors $\hat{\mathbf{n}}_i$ and $\hat{\mathbf{n}}_j$) which share an edge in the mesh experience a harmonic dihedral potential:

$$U_{\rm bend}(\Phi) = \kappa_{\rm dih}(1 - \cos\Phi), \tag{12}$$

where $\cos\Phi = \hat{\mathbf{n}}_i \cdot \hat{\mathbf{n}}_j$. $\kappa_{\rm dih}$ is the vesicle rigidity and this endows the membrane with a bending modulus given by $\kappa = \sqrt{3}\kappa_{\rm dih}/2$.[14]

We represent active filaments as semiflexible bead-spring polymers consisting of $L = 100$ beads of diameter $\sigma$, joined in a linear chain. Neighboring beads are connected by "tethers" similar to those mentioned above. For the active filaments $l_{\rm min} = 0.25\sigma$, $l_{\rm max} = 0.75\sigma$, $l_{c0} = l_{c1} = 0.5\sigma$. The strength of the tether interaction is set to $b = 1000k_{\rm B}T\sigma$

Deviations of adjacent bonds $\hat{\mathbf{b}}_i$ and $\hat{\mathbf{b}}_j$ from a parallel orientation are penalized with a harmonic angle potential:

$$U_{\text{angle}}(\Theta) = \kappa_{\text{fil}}(\Theta)^2, \tag{13}$$

where $\cos\Theta = \hat{\mathbf{b}}_i \cdot \hat{\mathbf{b}}_j$ and $\kappa_{\text{fil}} = 1000 k_{\text{B}}T$. Our simulations used 40 filaments.

Excluded volume interactions between non-bonded monomers are implemented with an expanded WCA potential:

$$U_{\text{WCA}} = \begin{cases} 4\epsilon\left[\left(\frac{\sigma}{r-\Delta}\right)^{12} - \left(\frac{\sigma}{r-\Delta}\right)^{6}\right], & r < r_c \\ 0, & r \geq r_c, \end{cases} \tag{14}$$

where $r_c = 2^{1/6}\sigma + \Delta$. For interactions between filament monomers, $\Delta = 0$. Filament-vesicle monomer interactions instead have $\Delta = \sigma_{\text{ves}}/2$, which prevents filaments from escaping the vesicle. Vesicle monomers do not experience excluded volume interactions with each other. We set $\epsilon = k_{\text{B}}T$.

Summing all such interactions yields the total potential energy of the system:

$$U_{\text{total}} = \sum_{\text{pairs}} U_{\text{WCA}}(r) + \sum_{\text{bonds}} U_{\text{tether}}(r) + \sum_{\text{angles}} U_{\text{angle}}(\Theta) + \sum_{\text{dihedrals}} U_{\text{bend}}(\Phi). \tag{15}$$

Our model evolves in time according to Langevin dynamics. Specifically, the equation of motion for bead $i$ within molecule $\alpha$ ($= 0$ for a vesicle bead, $> 0$ for a filament bead) is:

$$m\frac{\partial^2 \mathbf{r}_i^\alpha}{\partial t^2} = -\frac{\partial U_{\text{total}}}{\partial \mathbf{r}_i^\alpha} - m\gamma\frac{\partial \mathbf{r}_i^\alpha}{\partial t} + (1-\delta_{\alpha,0}) f_a \xi(t) \frac{\mathbf{r}_{i+1}^\alpha - \mathbf{r}_{i-1}^\alpha}{|\mathbf{r}_{i+1}^\alpha - \mathbf{r}_{i-1}^\alpha|} + \boldsymbol{\eta}_i^{T,\alpha}(t), \tag{16}$$

where $m = 1$ is the mass, which is identical for all beads. In this equation, the first term on the right hand side is the conservative force due to the potential energy; the second term is a friction force with damping coefficient $\gamma = 0.1\tau$ (where $\tau = (m\sigma^2/\epsilon)^{1/2}$ sets the system's timescale); the third term is the active force (or *activity*); and the fourth term is a Gaussian white noise representing thermal fluctuations. The active force acts tangent to the filament beads with a strength $f_a$ and reverses direction (modelling an active nematic) with a characteristic time $\tau_{\text{rev}}$. This reversal is accomplished via the Bernoulli process $\xi(t)$ which switches from $+1$ to $-1$ (or vice versa) with mean time $\tau_{\text{rev}} = \tau$. The thermal force $\boldsymbol{\eta}^T(t)$ has mean zero and variance satisfying the fluctuation-dissipation theorem:

$$\left\langle \boldsymbol{\eta}_i^{T,\alpha}(t) \cdot \boldsymbol{\eta}_j^{T,\beta}(t') \right\rangle = 6\gamma k_{\text{B}}T\delta_{ij}\delta_{\alpha\beta}\delta(t-t'). \tag{17}$$

Equation 16 is integrated in time using the velocity Verlet algorithm[1] with a timestep $dt = 10^{-3}\tau$. All of our simulations are performed with a custom version of the LAMMPS software package,[15] the source code for which is available on github: https://github.com/mattsep/lammps/tree/hagan-group.

To compute observables of interest, we first "equilibrate" our systems. During the initial 1 percent of the simulation, filament orientations and positions are randomised within the vesicle. After the initialisation, the system is allowed to run for $1 \times 10^4\tau$ before measuring observables during the "production" run. Observables are computed by averaging over $10^5$ configurations, sampled every $10\tau$ from "production" trajectories of length $\approx 7\times10^4\tau$. Averages are also taken over a maximum of 48 independent trajectories for each set of parameters.

One key observable is the force that the active filaments exert on the vesicle via excluded volume interactions (aka *interaction force.*) We defined the interaction force $\eta_i^a$ acting on vesicle bead $i$ as follows:

$$\eta_i^a = \sum_j \hat{\mathbf{n}}_i \cdot \mathbf{f}_{ij}^{\mathrm{WCA}}(r_{ij}), \tag{18}$$

where the sum runs over all filament beads $j$, $\hat{\mathbf{n}}_i$ is a unit vector normal to the vesicle at bead $i$, $r_{ij}$ is the distance between vesicle bead $i$ and filament bead $j$, and $\mathbf{f}_{ij}^{\mathrm{WCA}}$ is the excluded-volume force between the vesicle and filament beads:

$$\mathbf{f}_{ij}^{\mathrm{WCA}}(r) = \begin{cases} \frac{24\epsilon}{\sigma}\left[2\left(\frac{\sigma}{r-\Delta}\right)^{13} - \left(\frac{\sigma}{r-\Delta}\right)^{7}\right]\hat{\mathbf{r}}_{ij}, & r < (\Delta + 2^{1/6}\sigma) \\ \mathbf{0}, & \text{else,} \end{cases} \tag{19}$$

where $\hat{\mathbf{r}}_{ij}$ is the unit vector pointing from bead $i$ to bead $j$. As for the radial position of the vesicle beads, the interaction force can also be decomposed into spherical harmonics. Changing the vesicle bead label from the discrete index $i$ to continuous polar and azimuthal angles $\theta$, $\phi$, the interaction force can be written:

$$\eta^a(\theta,\phi) = \sum_{l=0}^{l_{\max}} \sum_{m=-l}^{m} \eta_{lm}^a Y_{lm}(\theta,\phi). \tag{20}$$

#### 4.3 Simulation results

##### 4.3.1 Measurements of forces on the vesicle validate assumptions made in analysis of experiments

**Filaments are confined at the vesicle boundary.** As seen in the experiments, the active filaments in the simulation spend most of their time at the vesicle boundary. This can be seen in Fig. 8a which shows a histogram of the radial position of active filaments in a "floppy" vesicle (with $\kappa_{\mathrm{dih}} = 50k_{\mathrm{B}}T$.) We expect this to happen because the persistence length of the active filaments(defined as the ratio of self-propulsion velocity and rotational diffusion coefficient of the filament) is much greater than the size of the vesicle. In other words, the filaments are strongly confined.

**Interaction forces are highly correlated with filament density.** We tested the assumption used in experiments that the interaction force $\eta_i^a$ due to the local density of filaments near a vesicle bead $i$ is a good proxy for the force

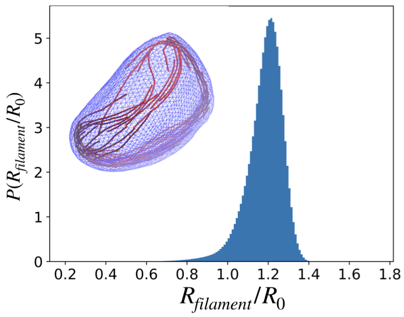


(a)

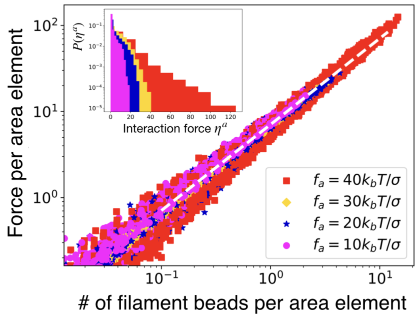


(b)

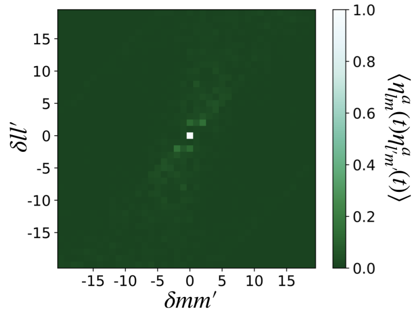


(c)

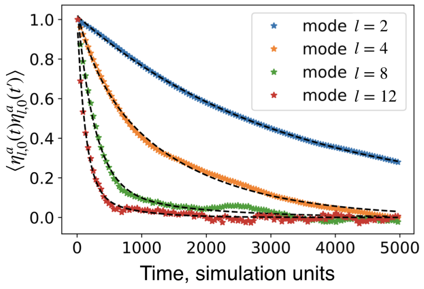


(d)

Supp.Fig. 8: Properties of the interaction force. The plots represent data for a vesicle with bending rigidity set to $\kappa_{\rm dih} = 50k_{\rm B}T$. (a) Histogram of filament positions showing that filaments are concentrated at the boundary of the vesicle. Inset - Snapshot of a typical vesicle configuration when the bending stiffness of the vesicle is set to $\kappa_{\rm dih} = 50k_{\rm B}T$. The tangential active force on the filaments is $f_a = 40k_{\rm B}T/\sigma$. (b) Scatter plot showing the proportionality between filament density and interaction force on the vesicle. The different colors and symbols represent vesicles containing filaments with different tangential active forces (or activities). The dashed white line is a linear function with slope of $\approx 7$. Inset - Histogram of forces exerted on the vesicle with filaments having different activities. The histogram is normalised such the area under the curves are is equal to 1. It can be inferred that force exerted on the vesicle is proportional to the local filament density for the range of activities. Filaments with a larger $f_a$ exert a larger force $\eta^a$ on the vesicle and the density of filaments interacting with the vesicle is higher. (c) Heat map of equal time force correlation functions versus differences in mode number, m-m' and l-l', for a vesicle with filaments that have a active force of $f_a = 40k_{\rm B}T/\sigma$. The color bar shows the value of the correlation function. It can be seen that the modes are only weakly correlated for $l \neq l'$ and $m \neq m'$. (d) Auto-correlation functions for the force on the vesicle for a few different modes $l$ (with $m = 0$.) These correlation functions are fit to an exponential function (Black dashed line represents the fit).

pushing outward on the vesicle. Fig. 8b shows a scatter plot of the average interaction force versus average local filament density. The averages were computed by binning the forces and densities over $\Delta\theta = 0.2$, $\Delta\phi = 0.4$ (corresponding to roughly 5 vesicle beads) and then averaging over timesteps. The local filament density, was calculated by counting the number of filament beads within a distance $2^{1/6}\sigma$ of the vesicle height centered within the solid angle bin. As can be seen, the interaction force is linearly correlated with the filament density over a range of activities (tangential active force on the filaments), supporting the experimental assumption. As a function of activity $f_a$, larger values of average interaction forces were observed for filaments with larger $f_a$. This can be seen in the histogram of the interaction force (inset to the Fig.8b). A greater number of filament beads could be found within our prescribed solid angle bin for filaments with greater active force on them which increased the total force on the vesicle. Note that the motion of active filaments was perpendicular to the surface normal of the vesicle and hence the force that the filaments exerted on the vesicle were largely due to extensile property of the active force. An increase in the active force therefore resulted in an increased number of filament bead interactions per vesicle bead which in turn resulted in larger values of the interaction force. The justification of this hypothesis can be understood in terms of the relationship between the average interaction force and average filament density in Fig.8b). The two quantities are linearly related with a slope is independent of $f_a$. This indicates that for a given number of filament beads the force on the vesicle is independent of the magnitude of the active force and only depends on the number of filament beads interacting with the vesicle.

**Force fluctuations in different spherical harmonic modes $(l, m)$ are approximately uncorrelated.** The equations used to model the experimental fluctuation spectra assumed that different modes of the interaction force $(\eta^a_{lm})$ are uncorrelated. We tested this assumption in our simulations by measuring the correlation function $\langle \eta^{a*}_{lm}(t)\eta^a_{l'm'}(t)\rangle$, where the average is taken over trajectories and over time within individual trajectories. As Fig. 8c shows, different spherical harmonic modes were largely uncorrelated, so that $\langle \eta^{a*}_{lm}(t)\eta^a_{l'm'}(t)\rangle \approx \langle |\eta^a_{lm}|^2\rangle \delta_{ll'}\delta_{mm'}$, as assumed in the analysis of experiments.

**Force fluctuations are exponentially correlated in time.** The autocorrelation function $\langle \eta^{a*}_{lm}(0)\eta^a_{lm}(t-t')\rangle$ was calculated with the average taken over all trajectories and initial times. We found that this function decays exponentially with time suggesting that $\langle \eta^{a*}_{lm}(0)\eta^a_{lm}(t-t')\rangle = \langle |\eta^a_{lm}|^2\rangle e^{-|t-t'|/]\tau_l}$. We inferred the corresponding mode relaxation timescale $\tau_l$ by measuring the time over which the correlation function decays by a value of $1/e$. Fig. 8d shows the force auto-correlation function for a few modes measured in floppy vesicles containing active filaments. The black dashed lines represent the exponential fits and their agreement with the simulation data is reasonable. This validates the assumption regarding the force fluctuations in the analysis of the experiments.

##### 4.3.2 Vesicle height fluctuations are enhanced by active forces

We analyzed vesicle height fluctuations by performing the spherical harmonic decomposition of Eq. 9 and measuring fluctuations of the spherical harmonic modes, $\langle|\delta f_{lm}|^2\rangle$, where $\delta f_{lm} = f_{lm} - \langle f_{lm}\rangle$. The triangular brackets indicates that the averaging of $|\delta f_{lm}|^2$ was done over different configurations in time and over all the different trajectories. To make a more direct comparison with experiments, we also computed fluctuation spectra projected onto a great circle (the "equator"):[11]

$$\langle|u_q|^2\rangle = \sum_{l=q}^{l_{\max}} \langle|f_{lq}|^2\rangle [Y_{lq}(\theta = \pi/2, \phi = 0)]^2, \tag{21}$$

where $q$ indexes Fourier modes about the equator.

**Active vesicle fluctuations violate a law of thermodynamic equilibrium.** In equilibrium the amplitude of fluctuations and the corresponding relaxation timescales for every mode are proportional. The proportionality constant depends on the mean vesicle radius, temperature, and effective mobility $\mu$ (inverse viscosity): $\langle|f_{lm}|^2\rangle = k_{\mathrm{B}}T\mu\tau_l/R_0^4$. As required by the fluctuation-dissipation theorem, the relaxation times of passive vesicle deformations are precisely determined by the magnitude of equilibrium height fluctuations. This is evident for an empty, passive vesicle as shown in Fig. 9a where the normalised magnitude of fluctuations $\langle|\delta f_{l0}|^2\rangle/\langle|\delta f_{20}|^2\rangle$ and the normalised relaxation timescale $\tau_{l0}/\tau_{20}$ are plotted as a function of mode number $l$. For vesicles with active filaments, this proportionality breaks down and the scaling laws for the two quantities are no longer the same. This is shown in Fig. 9b where the two are measured for a vesicle with active filaments having a force of $f_a = 40k_{\mathrm{B}}T/\sigma$. The relaxation timescale no longer scales according to the equilibrium expectation, suggesting that instead the force due to the filaments dictates the timescale of vesicle relaxation.

**The height fluctuations of vesicles increase monotonically with increasing active force on the filament.** The height fluctuations measured in 3D and summed up along a great circle along the vesicle allow us to make comparison with the experimental measurements of the same quantity. Fig. 9c and Fig. 9d show the height fluctuation spectrum as a function of mode number $q$ with different colors and symbols representing filaments with different active force on them. The values at different mode numbers are normalised by the height fluctuations of vesicle containing passive filaments at that mode number. This normalisation is done in order to compare Fig. 9c which is for a vesicle with bending rigidity $\kappa_{dih} = 5000k_{\mathrm{B}}T$(stiff) and Fig. 9d; for a vesicle with bending rigidity $\kappa_{dih} = 50k_{\mathrm{B}}T$(floppy). It can be seen that the magnitude of fluctuations is larger for vesicles with filaments having larger active force $f_a$. This is in agreement with the measurements from the experiments that show the enhancement of fluctuations in active vesicles compared to passive ones. The enhancement was larger in floppy vesicles, especially for the short wavelength

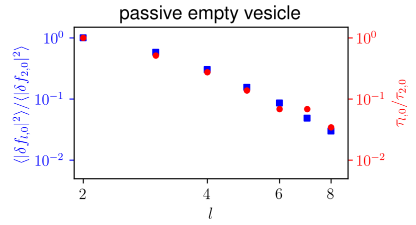


(a)

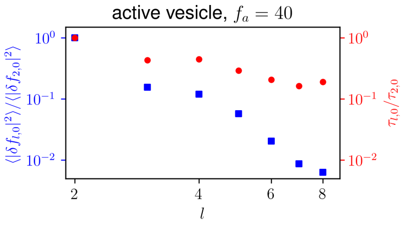


(b)

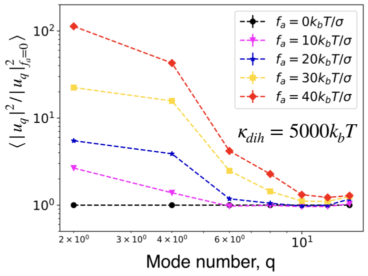


(c)

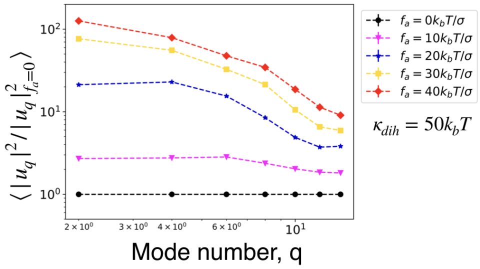


(d)

Supp.Fig. 9: Fluctuations of vesicles. (a), (b) The fluctuation spectrum for the spherical harmonic modes of the height function and the corresponding relaxation timescales as a function of mode number $l$. For an empty vesicle, these quantities are proportional as required by the fluctuation dissipation theorem (plot (a)). For a vesicle containing filaments with active force (plot (b)), the quantities are no longer proportional as the relaxation timescales of the modes decouple from the vesicles' material properties. (c), (d) The fluctuation spectrum for harmonic modes of the height functions summed up along a great circle on the vesicle. These are normalised by the the fluctuation spectrum of a vesicle containing passive filaments. As a function of active force $f_a$ on the filaments, it can be seen that the amplitude of fluctuations increases monotonically which is consistent with the experiments. For vesicles with different bending stiffness $5000k_{\mathrm{B}}T$ (on the left) and $50k_{\mathrm{B}}T$(on the right) we can see that the enhancement in fluctuations of short wavelength modes is larger for floppy vesicles for a given $f_a$. The short-wavelength modes of stiff vesicles are not enhanced because filaments do not impart enough force to access these high-energy vesicle configurations.

modes.

**Only the longest-wavelength modes of stiff vesicles showed enhanced fluctuations with increasing activity.** On comparing Fig. 9c and Fig. 9d, we can see that for very stiff vesicles ($\kappa_{\rm dih} = 5000 k_{\rm B}T/\sigma$,) active forces are unable to excite short-wavelength (high-$l$) vesicle modes. This underscores the important role that vesicle deformability plays in the experiments.

###### 4.3.3 Deformable vesicle relaxation times followed those of the interaction forces

The height auto-correlation function $\langle f^*_{lm}(0) f_{lm}(t-t')\rangle$ was calculated with the average taken over all trajectories and initial times. We found that this function decays as a double-exponential. The two timescales corresponding to the double exponential were attributed to the elastic relaxation of the vesicle (the longer time) and the relaxation of the vesicle due to dynamics of the interaction force (the shorter time). The results presented are with the shorter of the two times.

**The height relaxation times of deformable active vesicles followed interaction force relaxation times.** In contrast to equilibrium vesicles, the relaxation times of vesicles containing active filaments did not scale with $l$ in the same manner as the fluctuations (see Fig. 9b.) Instead, the vesicle relaxation times closely follow the relaxation times of the interaction force. Fig. 10b and Fig. 10d are scatter plots of the height relaxation timescales and interaction force relaxation timescales for different modes $l$. Fig. 10b shows modes $l < 6$ for vesicles with different bending rigidity (red circle: stiff vesicle with $\kappa_{\rm dih} = 5000 k_{\rm B}T$, blue square: floppy vesicle with $\kappa_{\rm dih} = 50 k_{\rm B}T$) both containing filaments with an active force of $f_a = 40 k_{\rm B}T/\sigma$. and Fig. 10d contains modes $6 \leq l \leq 15$. The black dashed line is a straight line with slope = 1. Fig. 10b indicates that the force and height relaxation timescale were equal for stiff and floppy vesicles alike for long wavelength modes. Fig. 10d however shows that this equality broke down for short wavelength modes, more so in the case of stiff vesicles. This is an important difference between floppy and stiff vesicles. The equality of the force and height relaxation timescales is still evident for floppy vesicles for short wavelength modes in Fig. 10d with only the point at $l = 15$ showing a significant deviation from the equality. In comparison, the equality of the two timescales for the stiff vesicle breaks down significantly for $l > 6$. The decoupling of the relaxation timescales suggests that the filament dyanmics is unable to excite short wavelength deformations in stiff vesicles.

**Role of vesicle deformability** It is evident from the comparison of height fluctuation spectrum and the correlation of height and force relaxation timescales that fluctuations of stiff vesicles due to interaction forces on them were unlike the fluctuations of floppy ones. The key differences are the enhancement of short wavelength modes (which were much smaller for stiff vesicles) and the decoupling of the height and force relaxation timescales (which is prominent for stiff vesicles). We attribute this inability of the filaments to excite the short wavelength modes to the fact that the energy required to excite them is quite high

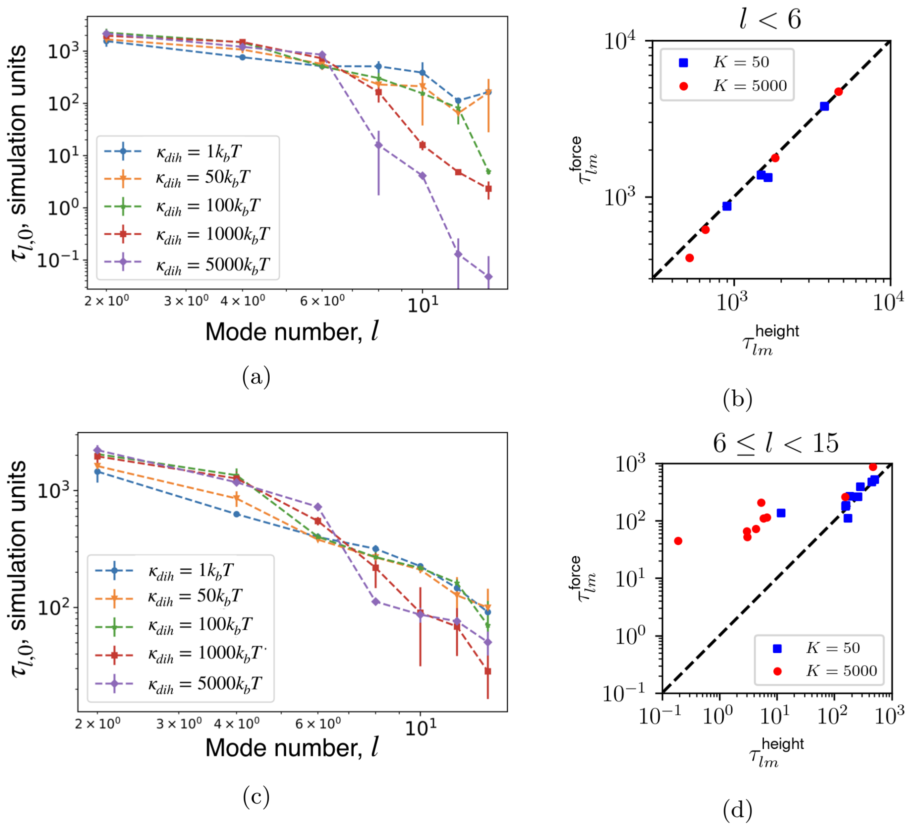


Supp.Fig. 10: Relaxation timescales for vesicles with different bending stiffness containing filaments with an active force of $f_a = 40k_BT/\sigma$. (a), (c) Relaxation timescales of harmonic modes for the height fluctuations(below) and force fluctuations(above) as a function of mode number $l$. It can be seen that the short-wavelength height fluctuations relax much faster for vesicles with higher bending stiffness (stiff vesicles). (b), (d) Scatter plots - relaxation timescales of height fluctuations vs force fluctuations. Panel (b) contains a scatter plot for points $l < 6$ and panel (d) contains points for $6 \leq l < 15$. The black dashed lines have slopes = 1 and serve to guide the eye. We can see that the two timescales are equal for long wavelength modes ($l < 6$). At short wavelengths, height relaxation times are no longer associated with force relaxation times for stiff vesicles.

for stiff vesicles. A comparison of the height relaxation timescales for vesicles with different vesicle rigidity as a function of mode number $l$ further substantiated our hypothesis. Fig. 10a shows this comparison and we can see that the vesicle height and interaction force relaxation timescales are comparable for long wavelength modes over the vesicle rigidity range we simulated. However, we see a sharp decline in the relaxation timescales for stiff vesicles at short wavelengths. We can compare these short wavelength height relaxations with the force relaxations in Fig. 10c. The decline in the force relaxation timescales is less significant for stiff vesicles at short wavelength.
